# *Scratcher*: An automated machine-vision tool for dissecting the neural basis of itch

**DOI:** 10.1101/2025.08.18.670778

**Authors:** Devangshu Nandi, Raghav Kaushik Ravi, Jagat Narayan Prajapati, Arnab Barik

## Abstract

Itch or pruritus invokes a specific reflexive and repetitive directed nocifensive behavioural response, known as scratching. Recent decades have revealed neural circuits that are involved in the sensory and affective-motivational aspects of itch-induced scratching. However, most of these studies relied on manual subjective methods of quantifying scratching in laboratory mice and rats. Recent advances in deep learning have opened avenues for the development of computational tools to analyze animal behaviour in a reliable and automated manner. Further, combined with optogenetic and chemogenetic strategies, these tools can accelerate our understanding of neural circuits underlying itch and scratching. To that end, we have developed *Scratcher*, a GUI-based computational tool based on a real-time object detection algorithm that allows semi-supervised automated analysis of scratching behaviour in mice in a computationally inexpensive manner. We recorded chloroquine-induced acute itch as it developed and determined the consequence of nail-trimming on acute-itch induced scratching with *Scratcher*. To probe the neural mechanisms underlying itch, we combined *Scratcher* with genetic circuit dissection using the Fos-TRAP mouse line. By targeting itch-activated neurons in the lateral parabrachial nucleus (LPBN) - a key brainstem hub for pruritic signal transmission - we demonstrated that LPBN activity modulates itch-evoked scratching. Together, we present a novel, easy-to-use computational tool to dissect molecular, cellular, and circuit mechanisms of itch and scratching.

## Introduction

The advent and application of machine learning and computer vision tools(Mathis et al., 2018; Pereira et al., 2022) have revolutionized behavioural neuroscience by making analysis of animal behaviours automated, semi-supervised, scalable, and reproducible. These computational approaches have accelerated data acquisition and interpretation and yielded mechanistic insights into complex behaviours, from locomotion to social interactions. Despite these advances, easily deployable and quantitative tools for measuring pruritic (itch-related) behaviours in rodents, which are the common laboratory models for studying itch, remain lacking. Computational tools that would require minimal exposure to programming languages and can be easily integrated with neural activity monitoring or closed-loop stimulation paradigms can help in a better understanding of the circuits underlying itch perception and processing. Importantly, despite its physiological and pathological relevance, the neural circuits underlying physiological and pathological itch remain poorly understood.

The sensation of itch is subjective, and a quantitative representation of its intensity is difficult; however, the behavioural output — scratching, can be objectively analyzed(Prajapati et al., 2024; Wimalasena et al., 2021). Over the last two decades, there have been efforts to automate the quantification of scratching behaviour in mice and rats. Hardware-dependent methods were developed to improve the specificity of scratch detection by using copper or aluminum rings that detect paw proximity to the nape of the neck(Elliott et al., 2000), a typical scratching target in rodents. These methods, however, often restrict free movement, potentially confounding behavioural readouts. Moreover, the use of magnetic or electric fields in these systems risks interfering with delicate neural recording techniques, thus limiting their utility in integrated systems neuroscience pipelines. In another development, force transducers placed on the floor of chambers were used to measure scratches in mice(Brash et al., 2005). In this inexpensive setup, a programmed microcontroller distinguished between scratching and grooming, and other non-specific movements simultaneously in multiple mice. Yet, they were limited in sensitivity and prone to both over- and under-representation of scratching events, especially when subtle or overlapping with other motor actions. More recently, an acoustic-based method was proposed wherein microphone recordings and signal processing algorithms were used to detect scratching sounds in freely moving animals(Elliott et al., 2017). While non-invasive and elegant in concept, this approach still suffers from limitations in resolving overlapping behaviors or disambiguating acoustic artifacts, ultimately limiting its robustness and specificity. Collectively, these approaches reflect sustained but cumbersome efforts to automate identification and quantification of scratching behavior, none of which have proven sufficiently accurate and generalizable across experimental contexts.

The emergence of deep learning-based pose estimation tools offered a promising new direction. Notably, Scratch-AID, an automatic itch detection framework, combined convolutional and recurrent neural networks (CNNs and RNNs) to classify scratching episodes(Yu et al., 2022). This approach marked a significant step forward by moving away from hardware-based proxies to directly model the behaviour from animal kinematics. However, despite its high precision, Scratch-AID requires substantial post-processing to distinguish scratching from kinematically similar behavior such as grooming. This tool introduces significant computational overhead and demands non-trivial pre-processing pipelines, which can compromise speed and scalability.

Given these methodological challenges, the need for a fast, accurate, and computationally efficient tool that can be deployed with minimal technical know-how remains unmet. Importantly, such a tool must be versatile enough to be adapted to analyze multiple animals simultaneously, work across different arenas, and integrate easily with neural recording or stimulation platforms - an increasingly common requirement in behavioural systems neuroscience research. To address these challenges, we developed *Scratcher*, an end-to-end, GUI-based behavioural analysis platform built on the YOLOv8 object detection framework(Redmon et al., 2016). Its computational efficiency and robustness across various experimental contexts make it ideally suited for long-term, high-throughput studies. With a focus on user-centric design and compatibility across experimental paradigms, *Scratcher* provides an accessible and scalable solution for the automated quantification of scratching behaviour, enabling broader adoption of objective and standardized itch analysis in rodent models. Notably, we employed *Scratcher* to study how the itch-sensitive ensemble of lateral parabrachial neurons participates in non-histaminergic acute, as well as spontaneous itch.(Mu et al., 2017). This approach indicates that *Scratcher* can be effectively deployed in itch circuit dissection studies.

## Results

### Development of *Scratcher*, a GUI-based deep learning tool to measure scratching in mice

We developed *Scratcher*, a computational tool to facilitate automated identification and quantification of scratching behaviour in an accurate and precise manner in mice. We developed the tool to provide the user with a suite of various quantitative measurements without compromising on computational cost in terms of time spent to obtain complete analyses. Further, the current iteration of *Scratcher* is specialized for detecting nape-directed scratching behaviour, since, in mechanistic studies, interrogating molecular, cellular, or circuit mechanisms of itch, pruritogens are typically injected or applied to the nape of the neck of mice. For acute itch, we injected 1% Chloroquine in the nape of wild-type CD1 mice (Figure 1A). We fabricated a behavioral setup (Figure 1B-C, see methods), an open-top box to enable unrestrained videography of mouse behavior with a matte dark background with uniform lighting of non-aversive intensity, to provide contrast to the coat color of the CD-1 wild-type strain of mice (Figure 1C). We leveraged the YOLOv8 object detection architecture to develop a deep learning-based pipeline to enable high-throughput, frame-accurate quantification of scratching behaviour from raw video data (Figure 1D-E). An object detection algorithm was selected over a classification-based approach(Oksuz et al., 2021) because it provides both spatial and categorical information, and not just detects whether a scratching event occurs, but also where in the frame it occurs. This localization capability is crucial for future extensions of the pipeline to closed-loop experimental paradigms, where real-time detection of scratch location can trigger targeted interventions(Mathis et al., 2018; Pereira et al., 2022; Schweihoff et al., 2021). Furthermore, object detection algorithms like YOLOv8 are significantly more computationally efficient during inference than frame-wise classification models(Hussain, 2024; Vasanthi and Mohan, 2024) which often require temporal context windows, optical flow, or full-frame analysis pipelines. Object detectors also allow for greater scalability(Dang et al., 2024; Hussain, 2024) as they can be adapted to detect multiple behaviours or features in parallel without restructuring the input format(Ge et al., 2024; Vasanthi and Mohan, 2024)

**Figure 1.**
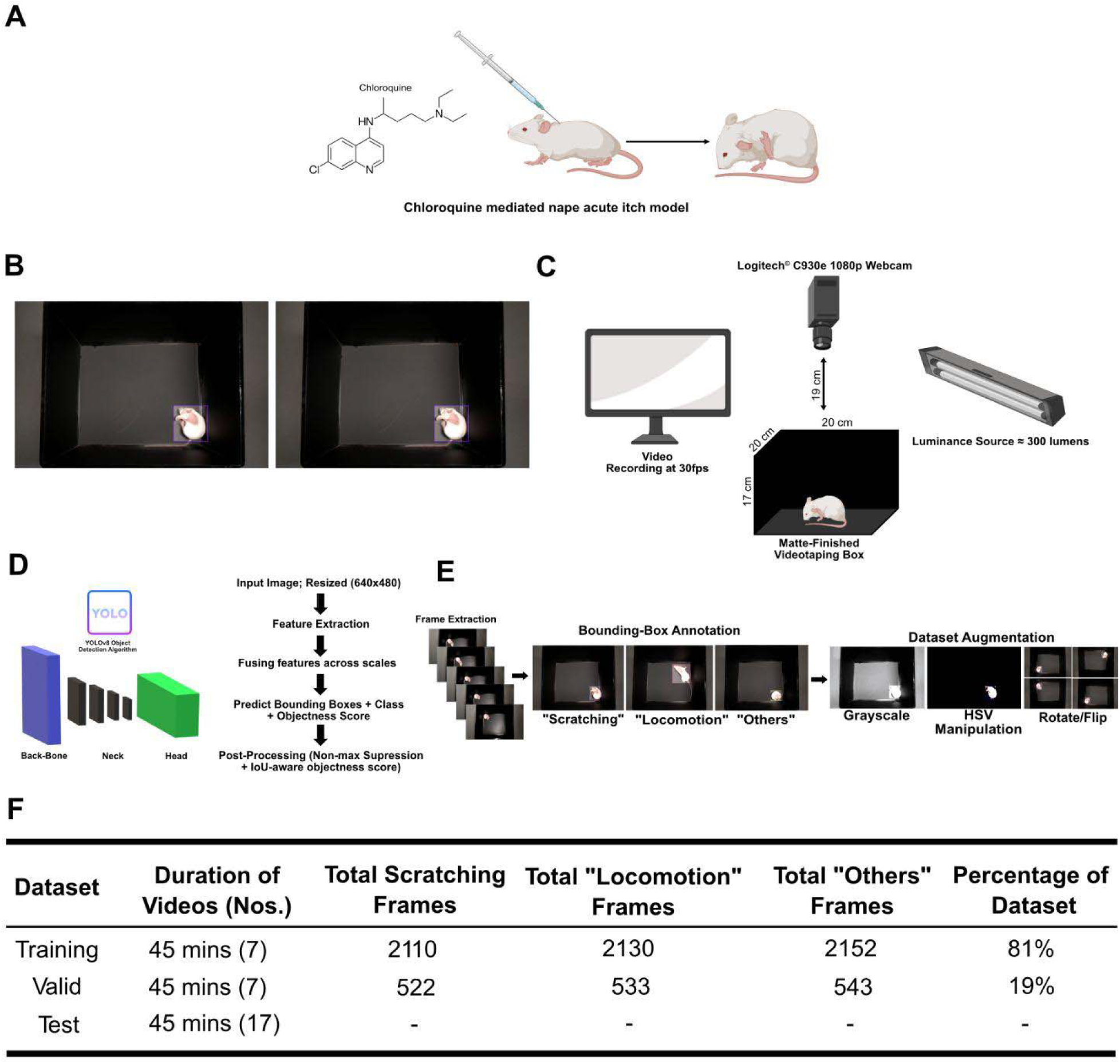
YOLOv8-Based Object Detection Pipeline employed in *Scratcher* for automated detection of scratching behaviours in various mice models of itch. (A) Schematic of the chloroquine-induced acute itch model, where intradermal injection of chloroquine at the nape of the neck evokes robust scratching behaviour. (B) Representative images of mice recorded from the top view in a standardized matte-black videotaping arena designed specifically for *Scratcher*. (C) Schematic of the video acquisition setup used in *Scratcher*. Mice were recorded under uniform lighting (≈ 300 lumens) using a Logitech 1080P business webcam mounted 19 cm above a custom matte-finished behaviour arena (20 × 17 cm), illuminated at 300 lumens. Videos were captured at 30 frames per second using the proprietary recording service LogiCapture 2.08.11. This precise setup is necessary to replicate for seamless compatibility with *Scratcher*’s GUI-based analysis pipeline. (D) Architecture of the YOLOv8-based object detection network used in *Scratcher*, highlighting key components including the backbone, neck, and detection head. (E) Workflow for dataset preparation and augmentation. Frame extraction is done from raw videos and manually annotated into three behaviour categories: “Scratching,” “Locomotion,” and “Others.” These are then augmented using grayscale conversion, HSV manipulation, and geometric transforms to enhance training diversity and prevent model overfitting. (F) Summary of the annotated dataset used to train, validate, and test the *Scratcher* detection algorithm. Each set contains 7 videos (∼45 minutes), with annotated frame counts per class and the relative proportion of the dataset across training, validation, and testing.

The workflow begins with standard video acquisition within a custom-designed behaviour box (for dimensions and other specifics see methods) optimized for visibility of scratching postures in freely moving mice (Figure 1C). Videos are parsed frame by frame, and each image is processed by a YOLOv8 neural network trained to detect and classify individual scratching events through bounding box localization. This bounding box approach(Segalin et al., 2021; Xu et al., 2025) provides precise spatial information about the scratch posture without altering the raw image data, thereby preserving the full anatomical context of the animal during behaviour. In contrast to other commonly used methods, such as background subtraction(Hu et al., 2023), which involve removing static elements from the video to isolate the mouse, bounding box detection is less computationally expensive(Sun et al., 2021) and avoids common pitfalls like partial removal or distortion of mouse body parts. Background subtraction can be particularly error-prone in behavioural settings(Wen et al., 2022; Zhao et al., 2019) where subtle movements and complex postures are critical, as it may inadvertently eliminate or modify important visual features necessary for accurate behaviour classification. By retaining the complete visual information and focusing on the relevant scratch regions, the bounding box method ensures that critical behavioural cues remain intact, enabling more reliable and robust detection across diverse experimental conditions(Wen et al., 2022; Zhao et al., 2019) YOLOv8 was selected for its high inference speed, real-time detection capability, and architectural advantages over previous versions, including a decoupled head structure, deeper CSPDarknet53 backbone, and robust anchor-free predictions(Redmon et al., 2016) (Figure 1D). These enhancements allowed the model to learn scratch-specific postures across diverse conditions with minimal overfitting, even when trained on a relatively small annotated dataset, while also reducing the spatial resolution and dimensionality of features in a manner that facilitates computation and increases overall processing speed, without compromising on accuracy. The final dataset consisted of a data split of 81% (6906 images)and 19% (1598 images) for training and validation, respectively (Figure 1F). This dataset consisted of videos from chloroquine-induced acute scratching, along with saline-injected rodent videos as controls. For testing various iterations of the models, we would test videos in which we were blind to the experimental condition. The entire open-source code and a full manual detailing the installation and usage of the GUI (Figure S1) can be found at its GitHub page (https://github.com/BarikLab-IISc/Scratcher).

### Evaluation of the model-training performance of *Scratcher*

After training the YOLOv8 model on an annotated dataset encompassing various postures associated with scratching, locomotion, and unrelated behaviours (“Others”), we assessed model performance using standard object detection benchmarks. Precision-recall analysis revealed a peak F1 score of 0.854 at a confidence threshold of 0.95, reflecting optimal balance between sensitivity and specificity for detecting behaviour-relevant bounding boxes (Figure 2A). Across the full prediction space, the model achieved a maximum average precision (mAP@0.5) of 0.967 (Figure 2B) and a corresponding average precision and recall of 0.97 and 0.99, respectively. (Figure 2C, 2D), indicating strong class separability and reliable retrieval of true scratching events. These metrics remained stable across a range of confidence thresholds, demonstrating consistent model behaviour across decision boundaries. Loss convergence was evident across training epochs, with box, class, and distributional focal losses (DFL) for both training and validation sets declining in a monotonic fashion (Figure 2E), consistent with stable gradient propagation and avoidance of overfitting. These trends were further supported by a steady increase in Mean Average Precision (mAP) and recall metrics across epochs, with class-wise spatial accuracy and confidence remaining robust across all behavioural categories (Figure 2G). The evolution of mAP was also tracked during training using two widely adopted evaluation standards: mAP@0.5, which considers predictions correct if their Intersection over Union (IoU) with ground truth exceeds 0.5, and the stricter mAP@[0.5:0.95], which averages mAP across multiple IoU thresholds (0.5 to 0.95, in 0.05 increments). As shown in the left panel (Figure 2H, left panel), mAP@0.5 rapidly increased during the initial epochs and plateaued at ∼0.94, indicating that the model learned to reliably localize and classify scratch events with moderate spatial tolerance. This suggests a strong ability to identify scratch frames with high sensitivity and low false positives under standard detection criteria. In contrast, the more stringent mAP@[0.5:0.95] metric (Figure 2H, right panel) showed a slower rise and reached a final value of ∼0.97. The relatively higher final value under the stricter metric demonstrates the model’s accurate predictions while exhibiting good spatial alignment with ground-truth annotations across a range of tolerances. A final confusion matrix constructed from the validation set confirmed near-perfect discrimination between the primary classes (“Itch”, “Locomotion”, and “Others”), with minimal confusion with the background class, further attesting to the model’s real-world applicability in noisy behavioural settings (Figure 2I). To operationalize the frame-wise annotations generated by the trained YOLOv8 model for downstream behavioural quantification, we implemented a structured post-processing pipeline (Figure 2J). This pipeline begins with the output files generated from the YOLOv8 model, which consist of frame-by-frame annotations of individual behaviors i.e., Itch, Locomotion, and Others, along with their associated confidence scores (Figure 2J, right) that proceeds to generate second-wise behavioural labels by aggregating frame-wise predictions using five distinct heuristics (details in the next section). We implemented and rigorously evaluated each of these heuristics, as each of them offers a different perspective on how high-frequency predictions (30 fps) can be transformed from noisy frame-level predictions into stable, second-level behavioral labels, making the data suitable for various downstream analyses.

**Figure 2.**
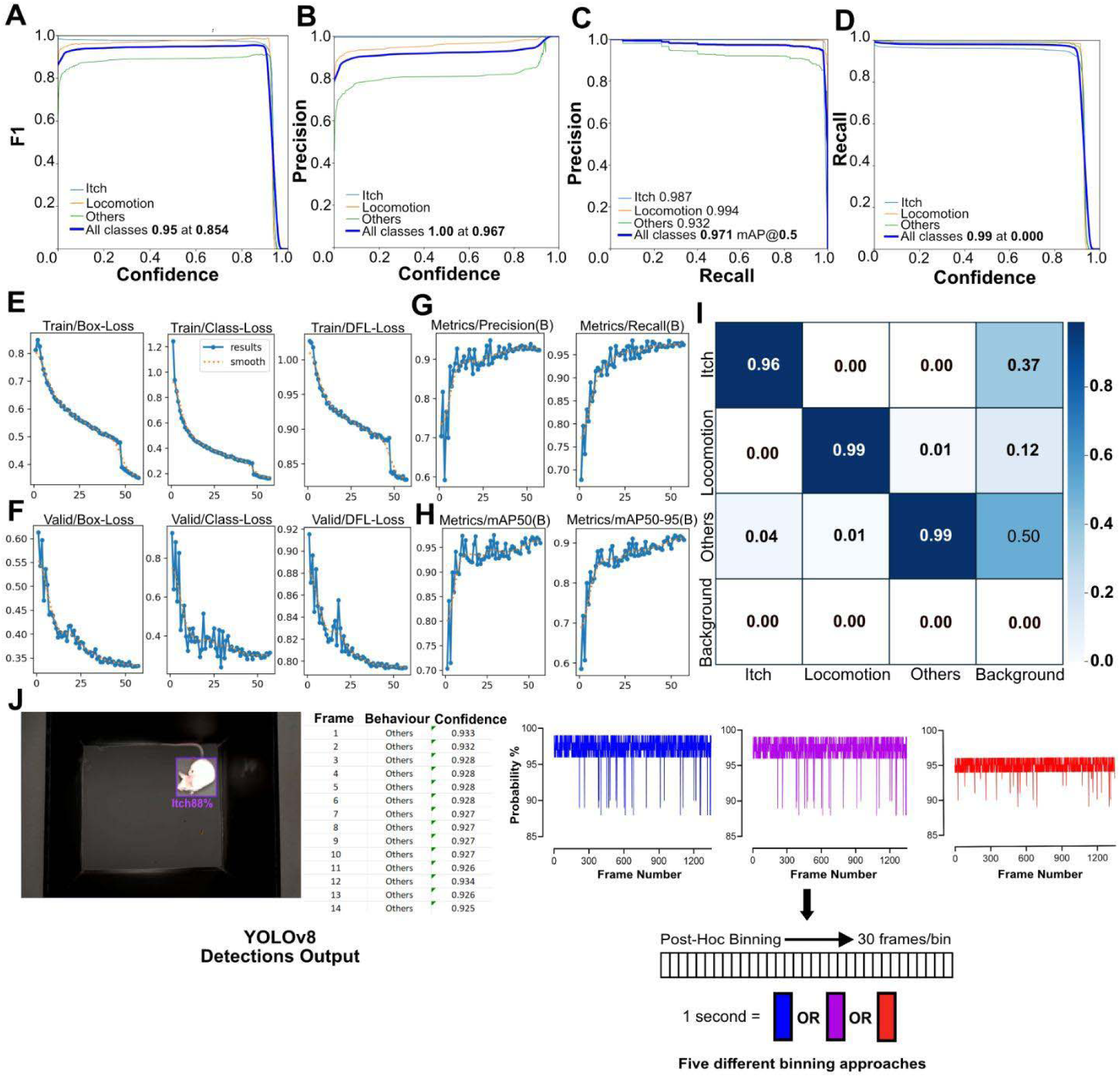
Quantitative evaluation and performance benchmarking of the YOLOv8 behaviour detection model and post-hoc binning strategies for obtaining second-wise behavioural classification. (A) Plot showing the F1 score as a function of confidence threshold for each behaviour class (“Itch,” “Locomotion,” “Others”) as well as the all-class average. (B) Precision vs. Confidence threshold curve across all behaviour classes. (C) Precision vs. Recall curve to benchmark the model’s capability to distinguish between behaviours in borderline confidence cases. (D) Recall vs. Confidence threshold curve for each behaviour class (E) Training loss curves over 50 epochs for bounding box regression loss (Box Loss), behaviour classification loss (Class Loss), and Distribution Focal Loss (DFL). (F) Validation loss curves for Box Loss, Class Loss, and DFL across 50 epochs. (G) Evolution of precision and recall metrics during training. (H) Mean Average Precision (mAP) curves during training, evaluated at IoU=0.5 (left) and at a stricter mAP@[0.5:0.95] standard (right). (I) Confusion matrix from the final YOLOv8 model output on the validation set. (J) Representative output from the trained YOLOv8 model. (Left) Single frame with predicted bounding box and class label (“Itch”) overlaid with confidence score. (Middle-left) Sample of raw frame-wise output showing predicted behaviour and corresponding confidence for each frame. (Middle-right) Probability traces of each class (“Itch,” “Locomotion,” “Others”) across all frames of a representative video. (Right) Post-hoc binning strategy for second-wise behavioural labeling, where 30 consecutive frames are grouped and labeled using one of five heuristic methods to derive robust time-resolved annotations for downstream analysis across the three behaviour classes (“Itch”, “Locomotion”, “Others”).

### A rule-based majority voting algorithm of post-hoc binning for Frame-to-Second behavioral aggregation using empirically derived tie-break rules

To convert frame-wise detections into behaviourally meaningful second-wise annotations, we developed a suite of post hoc aggregation strategies aimed at resolving the temporal complexity inherent in scratching behaviour. Unlike binary or posture-based motor acts, rodent scratching consists of a sequence of temporally extended yet phenotypically distinct subcomponents: (1) paw lift toward the nape (start), (2) rhythmic hind paw contact with the skin (scratching bout), (3) paw licking at the end of scratching, and (4) paw withdrawal (end). Although all four components comprise the canonical scratching sequence we are aware of, not all are equally suitable for classification in automated pipelines. For example, paw licking often occurs outside of itch-related contexts, and its appearance in isolation may not reliably indicate a scratching bout. In contrast, the scratching bout itself-characterized by rapid, repetitive, nape-directed hindlimb movements-is both temporally constrained and phenotypically distinctive, making it a robust unit for computer vision-based detection. Given the sub-second dynamics of these transitions and the need to deliver user-oriented outputs at behaviourally relevant timescales, we framed the binning task as a structured aggregation problem: the goal being to translate 30 frames of frame-wise predictions (at 30 fps) into a single second-wise behavioural label. This approach allowed the model to preserve the higher-resolution signature of scratching sequences while discarding ambiguous or non-specific micro-behaviours that often accompany scratching but are not an accurate diagnostic on their own. This approach also ensures that the resulting output reflects the user’s primary interest, i.e., nape-directed scratching, without being confused by grooming-like actions such as paw licking, which may have high visual overlap but completely different ethological relevance.(Shimada and LaMotte, 2008; Smolinsky et al., 2009; Wimalasena et al., 2021)

In the binning step, we first implemented a majority-voting heuristic applied over non-overlapping 30-frame windows, assigning to each second the most frequent behaviour observed within that window. A set of empirically derived tie-breaking rules was introduced to resolve ambiguous cases, prioritizing “Itch” in situations where licking or mixed behaviours co-occurred, based on manual inspection of over 40 hours of annotated recordings (Figure 3A). The resulting second-wise prediction series showed strong alignment with ground-truth annotations, as indicated exhaustively by several metrics. The confusion matrix revealed minimal misclassification between “Itch” and “Others” (Figure 3B), while time-aligned plots of prediction correctness further confirmed consistent performance across the session (Figure 3C, D). Quantitatively, the approach yielded an accuracy of 93.0%, precision of 94.3%, recall of 93.0%, and an F1 score of 93.3% (Figure 3E), validating the robustness of the rule set under noisy behavioural conditions. Distribution histograms showed strong agreement between predicted and actual class proportions (Figure 3E), including rare tie-label classes such as “Locomotion, Others” (Figure 3F). Precision-recall metrics remained high across individual classes (Figure 3G), and correlation analysis yielded an R² of 0.96, indicating strong agreement in cumulative behaviour counts (Figure 3H). Complementing this, a Bland–Altman plot was used to quantify agreement between the predicted and ground-truth scratch durations across timepoints (Figure 3I). This method, widely used in clinical and pharmacological research to assess bias and limits of agreement between measurement tools(Doğan, 2018; Ludbrook, 2010), revealed that most predictions fell within narrow agreement bands, suggesting negligible systematic bias. This indicates that our tool does not systematically overestimate or underestimate scratching duration and can be trusted for accurate behavioural quantification in experimental settings that have fundamental, as well as translational implications. To comprehensively assess the classification precision of Scratcher, we quantified the false negative rate (FNR) and false positive rate (FPR) for each behavioural class (Figure 3J). Across all annotated behaviours, Scratcher maintained a low FNR of approximately 0.14 and an even lower FPR of around 0.04, underscoring its robust ability to detect true behavioural events while minimizing false predictions. “Itch” events, in particular, were detected with high accuracy, with minimal omission or over-representation. The color-coded bar plots - blue for FNR and red for FPR - allow for intuitive visual inspection and demonstrate consistently low error rates across all behaviour types. These results highlight the tool’s strong reliability and reinforce its suitability for high-throughput, automated scratching behavioural analysis.

**Figure 3.**
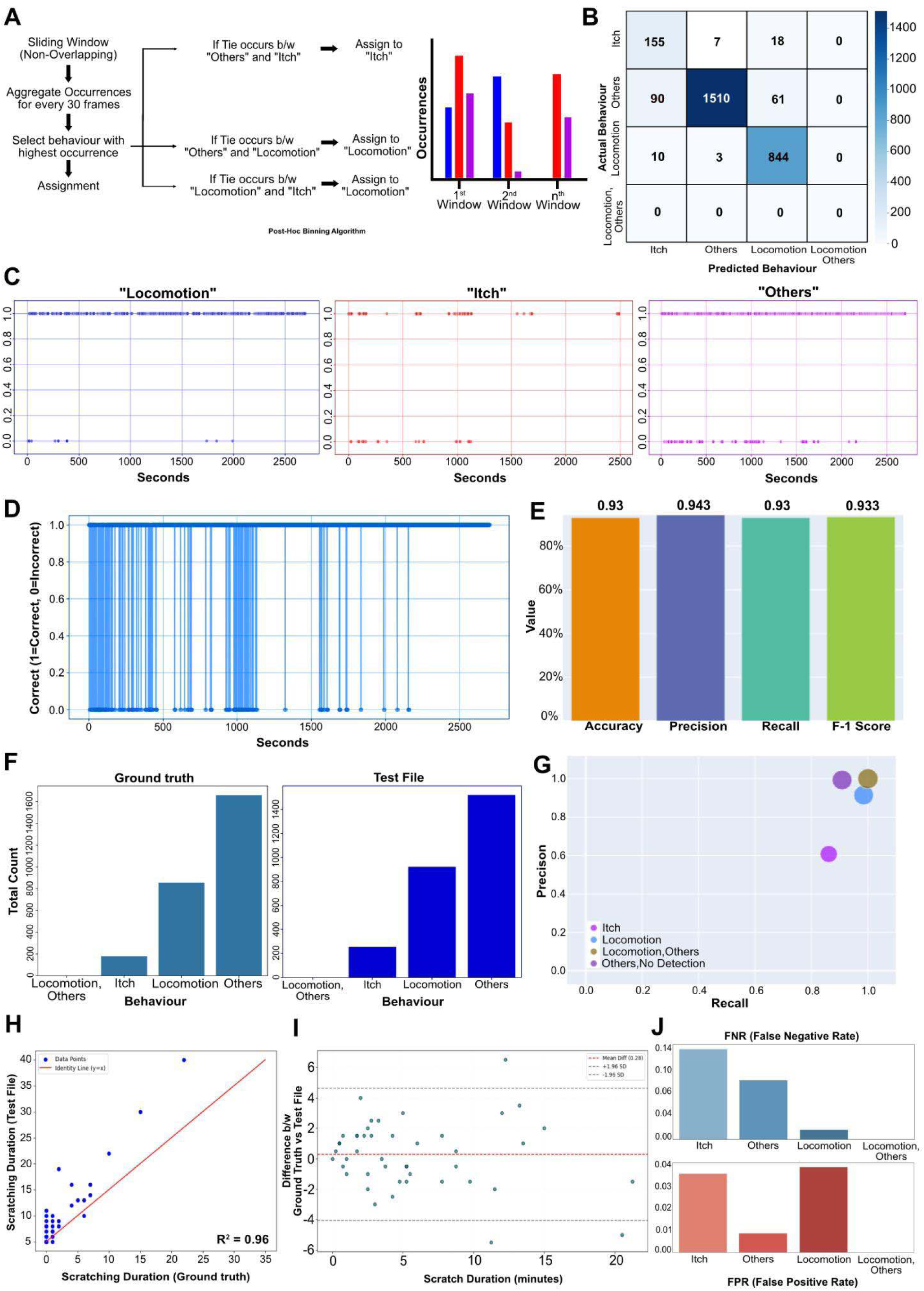
A rule-based majority voting algorithm of post-hoc binning for Frame-to-Second behavioural aggregation using empirically-derived tie-break rules. (A) Heuristic algorithm schematic. A step-wise schematic of the post-hoc binning process is shown. Within each 30-frame window (1 second), the behaviour with the highest frequency is assigned. In cases of ties between “Itch,” “Locomotion,” and “Others,” predefined rules prioritize assignment to “Itch” or “Locomotion” depending on the tie type. A representative bar plot of occurrences across three sample windows is included. (B) Confusion matrix comparing human-annotated ground truth with predicted labels. Rows represent true labels and columns represent predicted ones. (C) Accuracy of the temporal distribution of individual classes. Scatter plot showing the correctness of “Locomotion”, “Itch”, and “Others” (left to right) behaviour predictions over time, with y = 1 indicating agreement with ground truth (correct) and y = 0 indicating disagreement (incorrect). Each point represents a time-stamped prediction. (D) Temporal correctness across the test file. A dot is plotted for each second to denote whether the predicted label matches the ground truth (1 = correct, 0 = incorrect). Deviations from 1 indicate errors, allowing inspection of the temporal misclassification structure. (E) Classification metrics. Standard evaluation metrics—accuracy, precision, recall, and F1-score (F) Comparison of class-wise behaviour distributions. (G) Per-class Precision vs. Recall. Precision and recall values are plotted for each behavioural class (H) Correlation of estimated itch durations.A scatter plot comparing total itch duration per session between ground truth and test file (I) Bland–Altman analysis. Agreement between ground truth and test file is visualized via a Bland–Altman plot showing the mean difference and limits of agreement. (J) Bar graph displaying the False Negative Rate (FNR) and False Positive Rate (FPR) of the automated behavioural prediction system compared to human-annotated ground truth labels. The FNR bar, shown in blue, quantifies the fraction of actual “Itch” seconds that the model failed to detect. The FPR bar, shown in red, quantifies the proportion of seconds where the model falsely predicted “Itch” in the absence of a corresponding ground truth label.

Additionally, in order to arrive at a robust and generalizable heuristic, we evaluated five approaches across the same dataset, using manual human annotations as the ground truth. The first approach, a non-overlapping 30-frame majority voting scheme(Ren et al., 2020) with empirically determined tie-breaking rules, emerged as the most accurate and computationally efficient strategy (Figure 3). However, to evidence that this outcome was not incidental or biased by specific design choices, we present the data on the application of the other four algorithms. A Bayesian inference-based (Argiento et al., 2017; Beyer et al., 2013; McNamara et al., 2006) heuristic (Figure S2) was implemented to compute posteriors over behaviours by combining class priors (estimated from the training data) with likelihoods derived from YOLOv8 detection frequencies. While conceptually elegant, this approach struggled to maintain precision-recall balance, especially for “Itch”, which was consistently underestimated (Figure S2D–F). The algorithm tended to over-prioritize “Locomotion”, likely due to its dominance in the training set, and its performance suffered from the strong class imbalance. Temporal consistency in label assignment was also poor (Figure S2C), and aggregate accuracy metrics were significantly lower than those achieved via the majority voting scheme. Next, we tested a Gaussian Hidden Markov Model (HMM) (Macdonald and Raubenheimer, 1995; Patterson et al., 2009; Whoriskey et al., 2017), reasoning that it might better capture temporal dependencies in behavioural transitions (Figure S3). However, this approach performed poorly across all metrics, yielding an F1 score of just 11.1% (Figure S3D). The inferred hidden states exhibited significant confusion, especially between “Locomotion” and “Others”, and failed to reliably capture even the dominant behavioural class at any time point (Figure S3B). Ground truth distributions were not preserved (Figure S3E), and the overall correctness of predictions across time remained flat and near zero (Figure S3C). These findings suggest that while HMMs may be suitable for structured tasks with clean state transitions, they are ill-equipped for noisy, ethologically rich behaviours like spontaneous scratching, which do not conform to tightly ordered state machines. A third approach used a simple overlapping sliding window, where every window of 30 frames was moved one frame at a time and assigned the modal behaviour (Figure S4). While this improved temporal resolution, it introduced excessive redundancy, and its accuracy remained moderate (F1 = 46.9%). Notably, it severely overestimated “Locomotion” while underrepresenting “Others” and inconsistently detected “Itch” (Figure S4E–F). Despite some improvements in label smoothness, it did not outperform the non-overlapping majority vote. The fourth strategy, which averaged predicted class probabilities within non-overlapping 30-frame windows and assigned the behaviour with the highest mean confidence, showed considerable promise (Figure S5). This method maintained high confidence in its predictions and achieved strong performance across standard metrics, including an F1 score of 81.5%, the second highest among all tested heuristics (Figure S5D). Importantly, its predicted distribution closely matched the ground truth, though “Others” remained underrepresented (Figure S5E). Unlike other heuristics, this method exhibited high temporal correctness and a lower variance in frame-to-second translation errors (Figure S5C). These characteristics make it a viable alternative to the existing algorithm implemented in *Scratcher* currently for experiments where high prediction confidence is prioritized. Taken together, these analyses not only justify our use of the majority-voting scheme in the main pipeline but also demonstrate the breadth of computational approaches available for post hoc behaviour aggregation.

### Leveraging *Scratcher* to analyze itch-induced scratching behavior in mice

Scratcher was next evaluated for its ability to accurately quantify scratching behaviour for long-duration recordings, a capability essential for assessing the effects of prolonged drug treatments or neuromodulatory interventions. Typically, in studies designed to understand neural mechanisms of itch, post-injection of pruritogens in the nape of the neck, or cheek, scratching frequency is measured for 30 minutes to an hour. The reason for the choice of the time period for which behavior is usually assayed depends on the usual period for which the pruritogen-evoked scratching lasts at a high intensity, and the optimized time for which an experimenter can visually count the number of scratches. We argued that being able to record for a longer time period may delineate aspects of scratching behavior not revealed by shorter recordings. For example, longer recordings may help determine if neural manipulations affect the time for which the effect of pruritogens lasts. Animals sensitive to pruritogens may scratch at a similar frequency but for prolonged periods of time compared to the control conditions. Thus, we tested whether the *Scratcher* would allow us to analyze scratching behavior post-application of chloroquine in the nape of the neck of wild-type CD1 mice for three hours. We found that, despite the extended duration and increased behavioural variability, Scratcher maintained consistent frame-wise annotation accuracy and generated interpretable summary outputs, including cumulative behaviour plots and per-animal comparisons over time (Figure 4B–E). Individual animal trajectories revealed rich inter-animal heterogeneity, with some mice exhibiting sustained scratching activity over the full session (e.g., M1 and M6), and others demonstrating minimal or tapering responses (e.g., M3–M5) (Figure 4D). Linear regression analysis of scratching time courses revealed consistent negative slopes across most animals (Figure 4E), suggesting a decline in behavioural intensity over time. These results demonstrate that *Scratcher* can handle high-throughput behavioural sessions over extended durations without degradation in accuracy, offering a powerful solution for chronic assays or longitudinal tracking.

**Figure 4.**
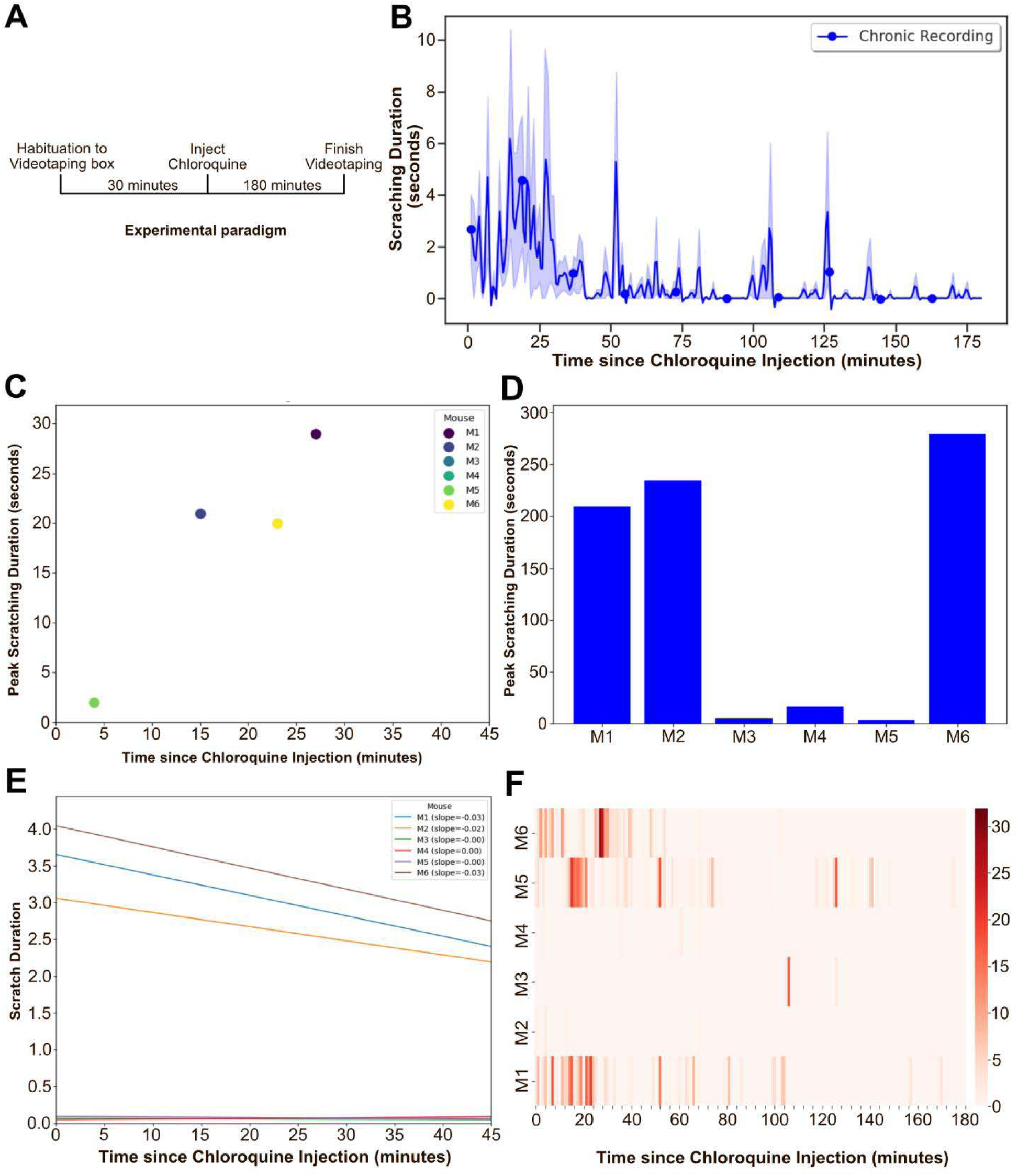
Scratcher enables high-resolution behavioural analysis during extended recordings. (A) Schematic of experimental protocol. Mice were habituated to the videotaping chamber for 30 minutes, followed by intradermal chloroquine injection. Scratching behaviour was then recorded for 180 minutes. (B) Scratching duration across a 3-hour session. Minute-wise average scratching durations across animals (n=6) overlaid on the individual data points (shaded area: SEM). (C) Time of peak scratching bout per mouse. Each dot represents the time (in seconds) post-injection at which each mouse exhibited its highest scratching duration. (D) Area under the curve (AUC). Bar plots showing cumulative scratching time (in seconds) across the session lasting for 3 hours. (E) Slope of scratching duration curve. Linear fits across time for all mice. (F) Heatmaps of scratching events. Each row represents a mouse, and each column a time bin (1 minute) across a full 3-hour session. Each row represents one mouse, and each column represents a 1-minute bin.

In the following experiments, we implemented *Scratcher* to ask if nail-clipping would alter itch-induced scratching patterns. Scratching an itch is known to bring relief; a lack of relief tends to increase the urge to scratch, and this increased urge is observed in chronic itch conditions. We hypothesized that clipping the nails of the mice would not provide them relief from chloroquine-induced itch by scratching, and thus, we should observe increased frequency of scratching bouts. We clipped the nails from the hind paws of mice and quantified chloroquine-induced scratching with the *Scratcher* before and after clipping (Figure 5A). We found that the scratching was increased after clipping, as expected (Figure 5B-E). Interestingly, the frequency of scratching was higher in the first 20 minutes in the mice after nail clipping (Figure 5B-E). Thus, when the pruritic effect of the chloroquine was the highest (time-post chloroquine administration), the nail clipping increased the scratching behavior compared to the unclipped condition. *Scratcher* analysis showed that the inter-animal variability was as expected (Figure 5C-E). Thus, with the help of an automated analytical pipeline, we show that the nail-clipping increases non-histaminergic pruritogen-induced scratching.

**Figure 5.**
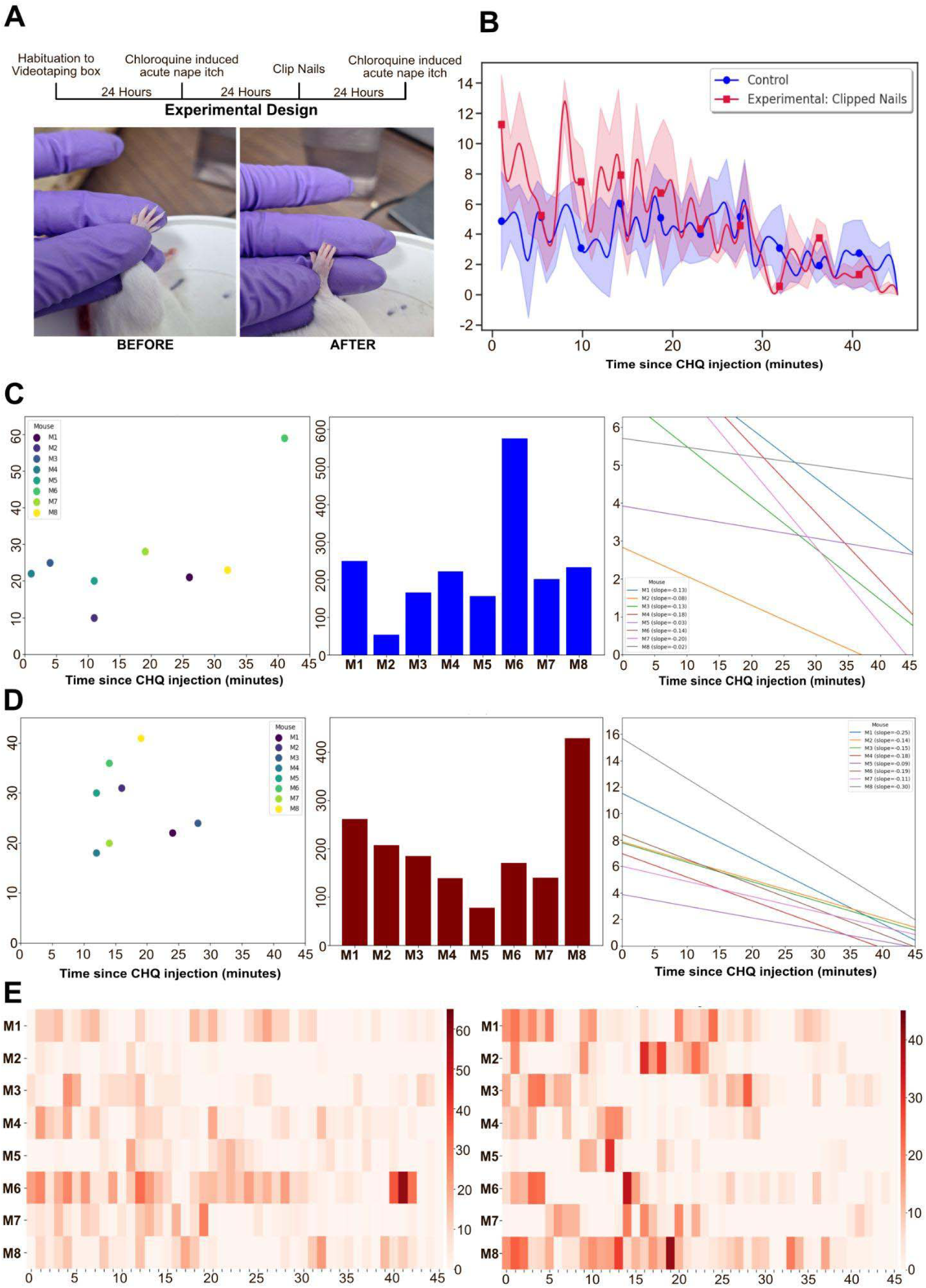
Scratcher enables precise quantification of scratching behaviour across experimental manipulations, such as nail clipping. (A) Experimental design and manipulation. Mice were habituated to the videotaping arena, followed by chloroquine (CHQ)-induced acute nape itch to establish baseline scratching. The next day, all forepaw nails were carefully clipped (image in set: BEFORE vs. AFTER), followed by another CHQ injection 24 hours later to assess changes in scratching behaviour due to mechanical alteration. (B) Group-wise time course of scratching bouts. Average scratch durations per minute across the session in control (pre-clipping, blue) versus experimental (post-clipping, red) groups. Shaded areas denote SEM. (n=8 mice) (C) Detailed scratch features for the control group (pre-clipping).Left: Peak scratch time per mouse. Middle: Total scratch duration per mouse in terms of Area under the curve (AUC) is relatively higher across all. Right: Linear slopes of scratching decay over time. Mice show individual variability, but overall maintain sustained scratching. (D) Detailed scratch features for the experimental group (post-nail clipping). (E) Heatmaps of scratching behaviour for individual mice before (left) and after (right) nail clipping. Each row represents a mouse; each column represents 1-minute bins.

### Effect of chemogenetic manipulation of itch-TRAP neurons in the LPBN with *Scratcher*

The LPBN is known as a central component of the pruritic ascending neural pathway (Mu et al., 2017; Prajapati et al., 2025; Ren et al., 2023), and we sought to understand the effects of chemogenetically activating the itch-sensitive neurons in the LPBN. To that end, we took advantage of the Fos-TRAP strategy, where a transgenic mouse strain (TRAP2) expresses 4-hydroxytamoxifen (4-OHT) sensitive Cre-recombinase under the immediate early-gene cFos promoter (DeNardo et al., 2019; Guenthner et al., 2013). In this transgenic strain, intraperitoneal administration of 4-OHT and simultaneous induction of neural activity induces expression of Cre in the cell population of interest (DeNardo et al., 2019; Onishi et al., 2024). We stereotaxically injected the AAV particles encoding Cre-dependent excitatory chemogenetic actuator tagged to mCherry fluorescent protein for visualization, hM3Dq-mCherry in the right LPBN of TRAP2, and induced the expression of the hM3Dq-mCherry in the LPBN itch-sensitive neurons by administering i.p. 4-OHT 120 minutes post-chloroquine injection in the nape of the neck of the mice (Figure 6A). 30 minutes after chloroquine administration is known to be sufficient for cFos induction, and 90 minutes is required for robust cFos-promoter driven gene expression(Guenthner et al., 2013; Samineni et al., 2021, 2019). Thus, we decided that 120 minutes after itch induction would be optimal for 4-OHT to drive Cre-expression in TRAP2 mice after pruritogen administration. We found that the itch-Trap drove robust hM3Dq-mCherry neurons in the LPBN, and i.p. DCZ administration induced cFos protein expression in the mCherry-positive neurons, implying successful chemogenetic tool induced neuronal activation (Figure 6B). We found that chemogenetic activation of the itch-sensitive LPBN neurons increased scratching (Figure 6C). Remarkably, the increased scratching was distributed across the 30 minutes for which the behavioral recordings were carried out (Figure 6D). Inter-animal variability was observed, which can be attributed to the differences in experimental conditions, levels of hM3Dq expression, and internal state of the mice (Figure 6E-H, Figure S6). Together, our findings indicate that targeted stimulation of the itch-responsive neural population in the LPBN is sufficient to drive increased nocifensive behavioral responses to pruritogens. Next, we wondered if the chemogenetic inactivation of the LPBN itch-TRAP neurons would alter the chloroquine-induced scratching in mice. To that end, we expressed the hM4Di-MCherry in the LPBN of TRAP2 mice through a Cre-dependent viral strategy and 4-OHT administration after a chloroquine-mediated itch assay (Figure 7A). We found that the chemogenetic inhibition of itch-TRAP neurons in the LPBN attenuated the chloroquine-induced scratching (Figure 7C). Importantly, the effect of chemogenetic inhibition on the scratching behavior was established within the first five minutes of the start of the assay, which was in turn started 10 minutes after i.p. DCZ administration (Figure 7D, Figure S6). Together, the LPBN itch-sensitive neurons are sufficient and necessary for the expression of scratching behavior induced by intra-dermal chloroquine in mice.

**Figure 6.**
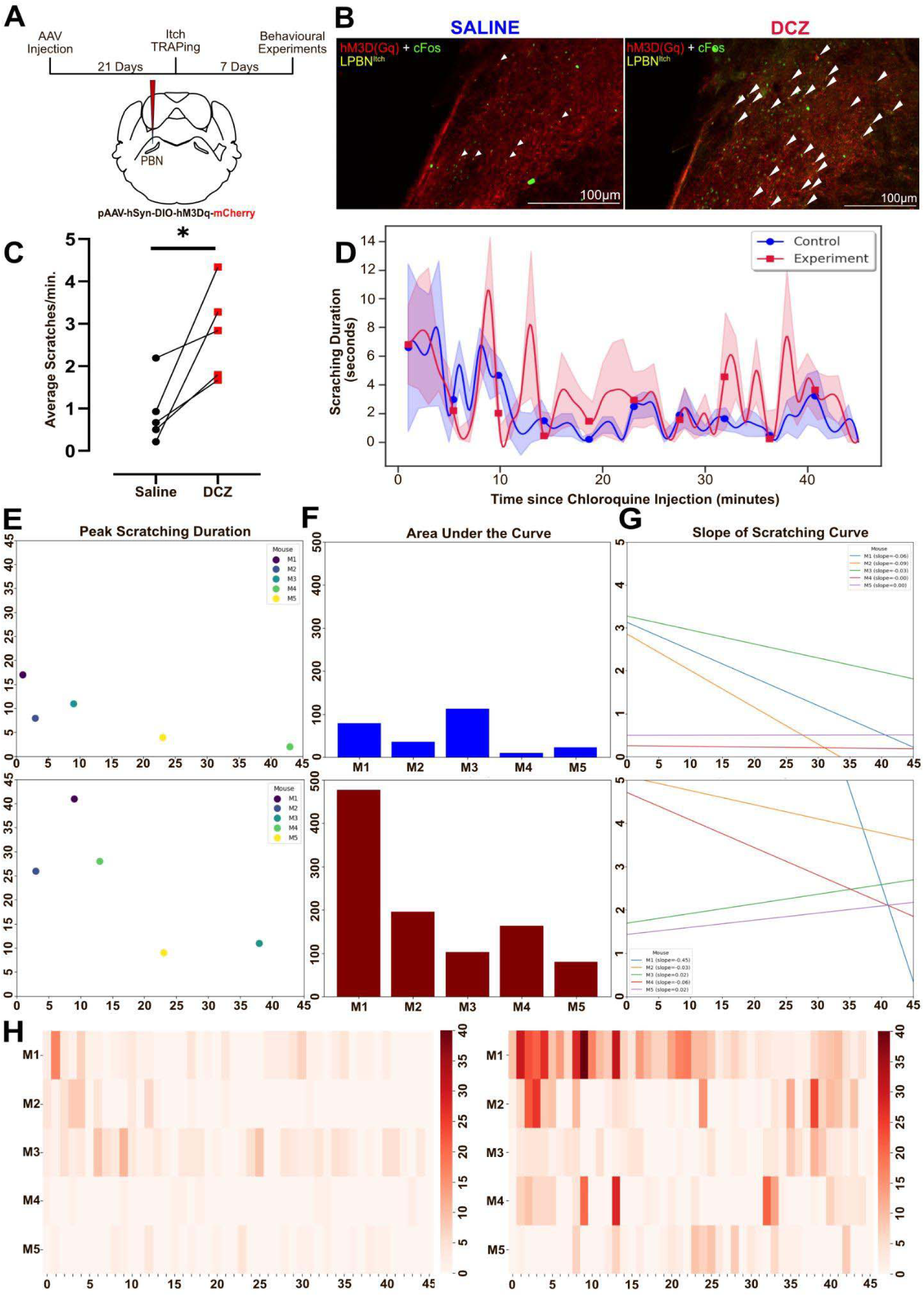
High-resolution profiling of scratching behaviour of scratching behaviour using Scratcher’s GUI-based analysis: Example output from user-supplied video demonstrating chemogenetic activation of Itch TRAPed neurons using multi-metric readouts. (A) Schematic of experimental design. AAV carrying Cre-dependent hM3D(Gq)-mCherry was injected into the parabrachial nucleus (PBN), and an itch TRAPing protocol was used to selectively express hM3D(Gq) in neurons active during chloroquine-induced itch. behavioural quantification was performed after saline injection and later after DREADD activation using DCZ (B) Histological validation of Itch TRAPed neurons. Representative sections show hM3D(Gq)-mCherry expression (red) and cFos (green) in the LPBN. (C) Average scratch bouts per minute across saline and DCZ conditions across all mice (n=5 mice, p < 0.05, paired t-test). Each dot represents one animal; red squares denote individual values in the DCZ condition. (D) Time-resolved scratch duration profile. Second-wise scratching duration over 45 minutes post-chloroquine injection across mice administered with saline(blue) and DCZ(red), respectively. Mean traces and shaded SEMs are plotted for all animals. (E) Peak scratching duration. Dot plots show the highest single-second scratching duration per mouse under saline (top) and DCZ (bottom) conditions. (F) Area under the curve (AUC). Bar plots showing cumulative scratching time (in seconds) across the session. (G) Slope of scratching duration curve. Linear fits across time show varying scratching trends for saline (top) and DCZ (bottom) conditions. (H) Heatmaps of scratching events. Saline (left) and DCZ (right), with each row representing a mouse, and each column a time bin (1 minute).

**Figure 7.**
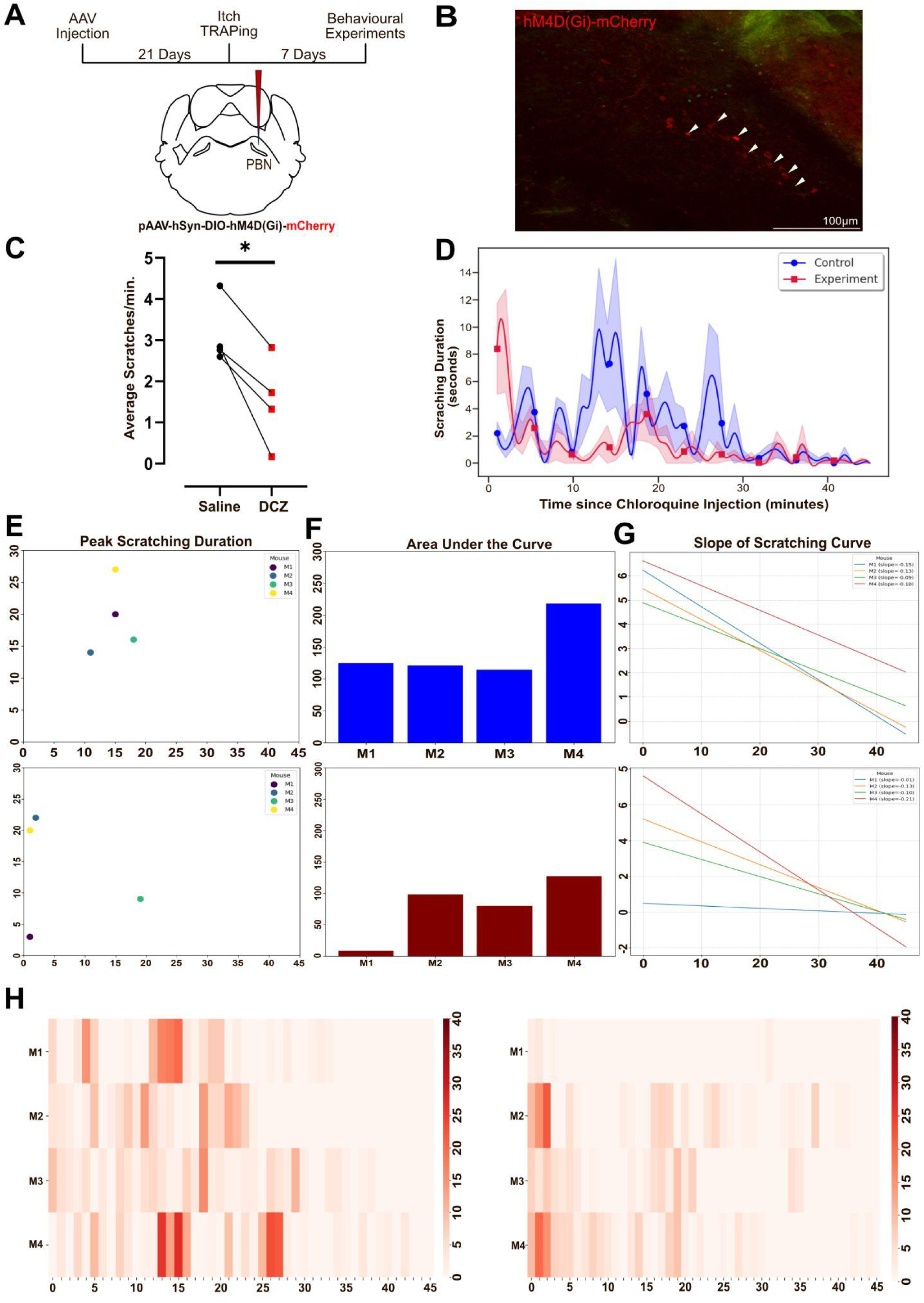
High-resolution profiling of scratching behaviour of scratching behaviour using Scratcher’s GUI-based analysis: Example output from user-supplied video demonstrating chemogenetic inactivation of Itch TRAPed neurons using multi-metric readouts. (A) Schematic of experimental design. AAV carrying Cre-dependent hM4Di-mCherry was injected into the parabrachial nucleus (PBN), and an itch TRAPing protocol was used to selectively express hM4Di in neurons active during chloroquine-induced itch. Behavioural quantification was performed before and after DREADD activation using DCZ. (B) Histological validation of Itch TRAPed neurons. Representative sections show hM4Di-mCherry expression (red) in the LPBN (C) Average scratch bouts per minute compared to saline controls across all mice (n=4 mice, p < 0.05, paired t-test). Each dot represents one animal; red squares denote individual values in the DCZ condition. (D) Time-resolved scratch duration profile. Second-wise scratching duration over 45 minutes post-chloroquine injection across DCZ (red) condition compared to saline controls (blue). Mean traces and shaded SEMs are plotted across animals. (E) Peak scratching duration. Dot plots show the highest single-second scratching duration per mouse under saline (top) and DCZ (bottom) conditions. (F) Area under the curve (AUC). Bar plots showing cumulative scratching time (in seconds) across saline (top) and DCZ (bottom) (G) Slope of scratching duration curve. Linear fits across time show varying scratching trends, saline (top) and DCZ administration (bottom). (H) Heatmaps of scratching events. Each row represents a mouse, and each column a time bin (1 minute). Saline (left) shows optimum scratching, whereas DCZ (right) reveals temporally sparse and minimal scratching episodes, captured with high temporal resolution by Scratcher.

### Simultaneous calcium imaging of the itch-TRAP neurons in the LPBN and *Scratcher*-mediated itch analysis

Finally, we demonstrate *Scratcher’s* compatibility with real-time systems neuroscience techniques by integrating it with the in vivo fiber photometry technique. In the fiber photometry technique, neural activity is measured indirectly from cell types of choice of behaving animals by expressing genetically encoded calcium sensors such as the GCaMP6s and digitizing the reflected light collected through a fiber optic cannula focused onto a CMOS camera with appropriate filters (Kielbinski and Bernacka, 2024; Simpson et al., 2024). In combination with *Scratcher,* fiber photometry recordings from a neural circuit of interest can inform us about the potential role of the circuit in the itch processing and scratching behavior. Here, we expressed the GCaMP6s in the itch-TRAP LPBN neurons following similar methods as above (Figure 6) and recorded the activity of the itch-TRAP neurons while the mice were injected with chloroquine in the nape of the neck. With the help of *Scratcher*, we extracted the timing of scratch bout onsets from the video data and aligned these to the calcium signal using a simple ΔF/F input file (Figure 8). This enabled automated generation of raster plots, bout-aligned heat maps, and peri-scratch ΔF/F traces (Figure 9B–E). Critically, comparison of these traces with manually aligned photometry data revealed close correspondence in both timing and shape of the average traces, affirming the temporal precision of Scratcher’s bout detection. Interestingly and in contradiction to previous studies on LPBN’s role in itch-induced scratching behaviors (Chen and Sun, 2020; Li et al., 2021; Mu et al., 2017; Pavlenko et al., 2025; Piyush Shah and Barik, 2022), we found that the LPBN itch-TRAP neural activity preceded the start of the scratching bouts. This seamless integration highlights *Scratcher*’s potential as a fully automated, high-throughput behavioural quantification interface for linking behaviour to neural dynamics without any need for manual adjustment or tuning.

**Figure 8.**
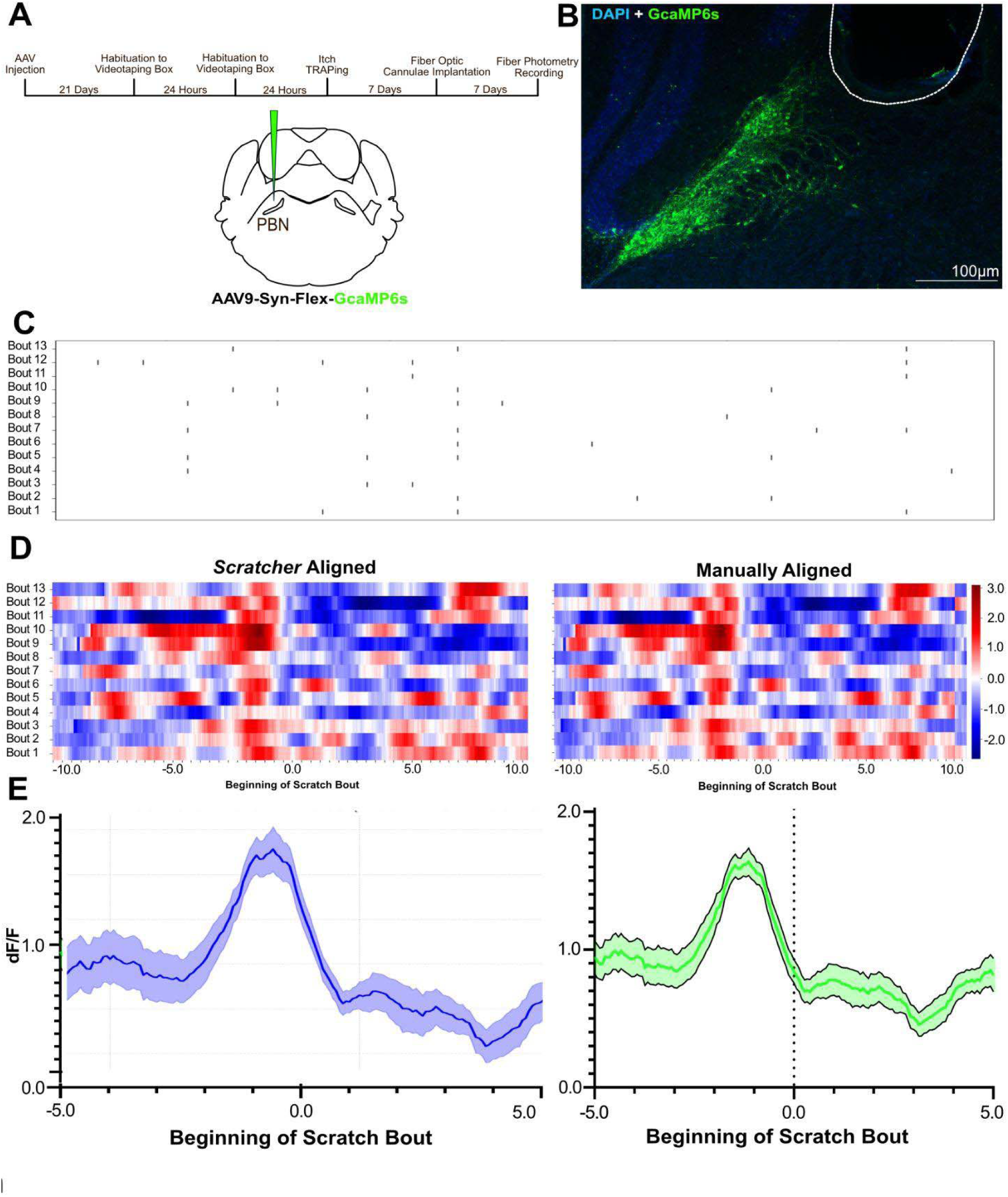
Scratcher integrates seamlessly with fiber photometry setups to automate behavioural alignment and neural activity analysis. (A) Schematic of the experimental timeline and viral targeting. Mice were injected with AAV9-Syn-Flex-GCaMP6s in the lateral parabrachial nucleus (LPBN), followed by habituation to the videotaping box, itch TRAPing, fiber optic cannulation, and performing fiber photometry recordings after injecting chloroquine to induce acute itch over the course of several weeks. (B) Histological validation of virus expression in the LPBN. Coronal section showing GCaMP6s expression (green) localized to the PBN, counterstained with DAPI (blue). Scale bar: 100 μm. (C) Automated peri-scratch event alignment by Scratcher. Top: Raster plot showing the time of start of individual scratching bouts across a 45-minute session. Bottom: Aligned fluorescence (ΔF/F) heatmap showing GCaMP6s activity 10 seconds before and after each scratching bout. Each row represents a single bout, while each column represents fluorescence values, automatically detected and aligned by Scratcher. (D) Averaged peri-bout GCaMP6s activity across all scratching events. Mean ΔF/F trace (green) shows a rise in LPBN activity leading up to a scratching bout, followed by a gradual return to baseline levels of fluorescence. The shaded region indicates SEM. Time zero denotes the onset of each scratching bout.

## Discussion

The study of itch has long been constrained by the lack of standardized, automated, scalable, and objective tools for quantifying scratching behavior in rodent models. To that end, we present *Scratcher*, a computationally efficient, accurate, and easy-to-deploy behavioral quantification tool for pruritic behavior. Importantly, we aimed to create a solution that could seamlessly integrate with experimental paradigms involving pharmacological, genetic, and circuit-level manipulations, while minimizing user-side programming or hardware requirements. Our approach, which is based on the YOLOv8 object detection framework and supported by an intuitive graphical user interface - provides a unified pipeline for the detection, aggregation, and analysis of scratching behavior from raw video data. We used *Scratcher* to demonstrate the ability to quantitate scratching behavior induced by a non-histaminergic pruritogen such as chloroquine. Importantly, we employed *Scratcher* to determine the effects of chemogenetic manipulation of the itch-responsive neurons in the LPBN, a critical node in the ascending neural pathways responsible for transmitting pruritic information from the dorsal spinal cord to the mid- and fore-brain structures (Mu et al., 2017; Prajapati et al., 2025, 2024). Further, we combined *Scratcher* with *in-vivo* calcium imaging to provide automated registration of itch-induced scratching behavior with neural activity. Together, we provided comprehensive evidence for the utility of *Scratcher* as a standardized analytical pipeline that helps uncover neural circuit mechanisms underlying scratching caused by pruritus.

Quantifying scratching through manual annotation, although informative, is subjective and time-intensive, making it poorly suited for high-throughput or long-duration studies(Le Bras, 2023; Sakamoto et al., 2022; Wimalasena et al., 2021). There have been several efforts to automate scratch behavioral analysis. Hardware-based approaches, which ranged from force sensors to proximity-based electrical systems, all have historically provided partial solutions but suffer from critical drawbacks, including mechanical interference, limited specificity, and incompatibility with systems neuroscience techniques(Chun et al., 2021; Deuis et al., 2017). Pose-estimation pipelines like Scratch-AID represent important progress, yet the need for extensive pre-processing, susceptibility to motion artifacts, and challenges in scaling across experimental conditions limit their usability for broad applications(Chen et al., 2022; Kobayashi et al., 2021; Sakamoto et al., 2022). In that note, *Scratcher* provides a generalizable, intuitive, and accurate application with GUI for itch studies.

Our results validate the design decisions that underpin Scratcher. First, the use of YOLOv8 allowed us to bypass many of the issues inherent to classification-based or background-subtraction-based methods (Figure 1-E). The one-shot detection architecture offered superior speed without compromising on accuracy, enabling real-time analysis that is well-suited for eventual closed-loop integration (Figure 1-E). Importantly, bounding box-based object detection provided both spatial and categorical behavioral information - allowing users not only to identify when scratching occurs, but where in the frame it happens (Figure1-F). This localization ability has immediate utility in multi-animal experiments and opens the door for future real-time manipulations guided by behavioral state.

To mitigate the challenges of translating high-frequency frame-level predictions into meaningful behavioral metrics, we systematically compared multiple second-wise binning heuristics (Figure 3, Figure S2-5). Among these, a non-overlapping majority vote strategy with rule-based corrections emerged as both robust and computationally efficient, achieving >93% precision, recall, and F1 scores (Figure 2A-D). The comparative evaluation of alternative heuristics - including Bayesian, Hidden Markov Model (HMM), confidence-based, and sliding window approaches - revealed the limitations of probabilistic methods in modeling the noisy and ethologically complex nature of scratching, while highlighting the importance of context-aware but interpretable aggregation strategies (Figure S2-5). Each heuristic reflects a different trade-off between temporal smoothness, class balance, interpretability, and computational cost. Importantly, our results highlight that no single approach is universally superior across all metrics. For instance, while Bayesian inference and HMMs offer principled probabilistic frameworks, they perform poorly in practice due to the noise and spontaneity of real-world scratching behaviour. The exhaustive comparison offered here provides a blueprint for users to select binning strategies suited to their specific experimental contexts. By validating our tool’s robustness across a spectrum of binning algorithms, we enable users to flexibly adapt the tool to a range of behavioural paradigms, including those that involve pharmacological, genetic, or environmental manipulations - or where such manipulations are absent and spontaneous behaviour is of primary interest.

Critically, Scratcher demonstrated strong generalizability across a range of experimental conditions. Whether applied to acute itch induced by chloroquine, or altered mechanical conditions introduced by nail clipping, the tool consistently captured behaviorally relevant changes. Its ability to sensitively and reliably quantify both enhancements and suppressions in scratching behavior was further demonstrated in chemogenetic experiments targeting Itch TRAPed neurons in the lateral parabrachial nucleus (LPBN). Not only could Scratcher detect modulation of opposing behavioural phenotypes following hM3Dq and hM4Di DREADD activation, it could also generate rich metrics such as slope analyses, peak scratching durations, area under the curve (AUC) and even latency to scratch (Figure S6) from the start of a session from simple video inputs, all via a user-friendly GUI. Moreover, we demonstrated the temporal precision of Scratcher in a systems neuroscience context by aligning bout onsets with *in vivo* calcium signals recorded from the LPBN. The high degree of temporal fidelity between automatically and manually annotated peri-scratch ΔF/F traces suggests that Scratcher can be a reliable companion to photometry workflows, and potentially extendable to closed-loop neural modulation frameworks.

Despite these strengths, several limitations remain. Currently, Scratcher is optimized for nape-directed scratching in singly housed mice under standard illumination and fixed camera positioning. While the tool can be retrained for other body sites or behaviors, this process still requires labeled datasets and may not generalize to more complex social settings without further adaptation. Accordingly, the bounding-box architecture was adopted as it is also inherently suited to multi-animal analysis. However, the current implementation does not perform individual tracking, which is essential for experiments involving more than one subject per frame. Integrating identity tracking into future versions could extend Scratcher’s capabilities to social paradigms, drug screening in group-housed mice, or genetically heterogeneous populations.

## Materials and methods

### Mouse line and treatments

Animal care and experimental procedures were performed following protocols approved by the CPCSEA at the Indian Institute of Science. TRAP2 (Fos2A-CreERT2) mice, stock number 030323, were purchased from Jackson Laboratory. For the rest of the studies, wild-type CD1 mice were used. The animals were housed at the Central Animal Facility under standard transgenic animal housing conditions in a 12-hour light-dark cycle with *ad libitum* access to food and water. Genotyping was performed according to the protocols (for iCre) of Jackson Laboratories. All mice used in the behavioural assays were between 8 and 12 weeks old. All the behaviours were done during the light cycle.

### Viral Vectors

Vector used and sources: pAAV5-hsyn-DIO-hM3D(Gq)-mCherry (Addgene, Catalog No. 44361, Titer: 1.8 × 10^13^ GC/ml), pAAV-h8yn-DIO-hM4D(Gi)-mCherry (Addgene, Catalog No. 44362, 2.5×10^13^ GC/mL), AAV9. syn.flex.GcaMP8s (Addgene, Catalog No. 162377, Titer: 2.7 × 10^13^ GC/ml),

### Stereotaxic injections

Mice were anesthetized with 2 % isoflurane/oxygen before and during the surgery. An incision was made to expose the skull, and subsequently, the skull was aligned to the horizontal plane. Craniotomy was performed at the marked point using a hand-held micro-drill (RWD). A Hamilton syringe (10 ul) with a glass pulled needle was used to infuse 300 nL of viral particles (1:1 in saline) at a rate of 100 nL/min. The following coordinates were used to introduce the virus: LPBN - AP: −5.34, ML: ±1.00, DV: −3.15.Post-hoc histological examination of each injected mouse was used to confirm that viral-mediated expression was restricted to the target nuclei.

### Itch TRAPing

4-Hydroxytamoxifen (4-OHT; Hello Bio, UK, Catalog No. H6040) was prepared by dissolving it in ethanol at a concentration of 20 mg/ml. The solution was subsequently aliquoted and stored at −40°C. 4-OHT was redissolved just before use and mixed with corn oil in a 1:1 ratio. 4-OHT (50 mg/kg body weight) was intraperitoneally administered to the mice, and 15 minutes later, the mice were subjected to chloroquine mediated acute itch in the nape. All the behavioural and anatomical studies were done one week after the Itch TRAPing.

### Videotaping Box

A rectangular acrylic box with dimensions identical to the homecage that the mice are housed in was designed to ensure naturalistic cage behaviour. The acrylic box was designed such that the interior of the box has a matte finish in order to ensure diffused luminance with no abnormally bright spots. This also ensures that no reflections arise that could interfere with the model’s detection or training. The dimensions of the box are as follows: 17cm (height) x 20cm (length) x 20 cm (breadth)(Indus Biosolutions, Bengaluru). The videotaping camera used was a Logitech C920e Business Webcam. The video recording software used was Logitech Capture 2.06.12 Software, using the following parameters during recording: Brightness 50, Contrast 50, Resolution 1920×1080, Frame rate 30 fps. An additional LED attachment with a moving arm was also installed for ambient lighting. The lighting was always maintained at approximately. 300 lumens are placed at a distance of 14 cm above the highest point of the videotaping box.

### Chloroquine-induced itch assay

The nape of the neck of mice was shaved 2–3 days before behavioural experimentation, and the mice were habituated in the behaviour room. On the day of experimentation, mice were individually placed in our custom videotaping box and habituated to it for 15 minutes. Chloroquine (375 µg/75 µl) was administered intradermally into the nape of the neck of the mice, and the subsequent scratching behaviour was recorded for 45 min (acute itch) or 3 hours, for acute and chronic recordings, respectively. The PC with the following specifications: Intel(R) Core(TM) i7-10700 CPU @ 2.90GHz, 64.0 GB RAM, 4GB NVIDIA GeForce GT 730, Windows 10 Education. For chemogenetic activation and inactivation experiments, DCZ (2 µg/kg of body weight) was administered intraperitoneally (i.p.) 15 min before chloroquine injection(Nagai et al., 2020). Hind leg-directed scratching of the nape was characterized as a scratch.

### Fiber optic cannula implantation

Fiber optic cannula from RWD (Ø1.25 mm Ceramic Ferrule, 300 μm Core, 0.39NA, L = 6 mm, Catalog No. R-FOC-BL300C-39NA) were implanted at AP: −5.34, ML: ±1.00, DV: −3.15 in the LPBN of the AAV-DIO-GCaMP8s itch TRAPed mice. Animals were allowed to recover for at least 1 week before performing behavioural tests. Successful labeling and fiber implantation were confirmed post hoc by staining for GFP for viral expression and injury caused by the fiber, respectively. Only animals with viral-mediated gene expression and fiber implantations at the intended locations, as observed in post hoc tests, were included in the analysis.

### Fiber photometry

A single-channel fiber photometry system from RWD (R810) was used to collect the data. The light from two light LEDs (410 and 470 nm) was passed through a fiber optic cable coupled to the cannula implanted in the mouse. Fluorescence emission was acquired through the same fiber optic cable onto a CMOS camera through a dichroic filter. The photometry data was analyzed using the RWD photometry software, and .csv files were generated. All trace graphs were plotted from .csv files using GraphPad Prism software version 8.

### Immunostaining and fluorescence microscopy

Mice were anesthetized with isoflurane and perfused transcardially with 1X Phosphate Buffered Saline (PBS) (Takara catalog No. T9181) and 4 % Paraformaldehyde (PFA) (Ted Pella, Inc. Catalog No. 18505). Harvested brains were further fixed in 4 % PFA overnight and subsequently transferred to 15 % and 30 % sucrose for serial dehydration. Brain tissues were placed in the Cryo-Embedding Compound (Ted Pella, Inc.) and frozen at −40°C. Subsequently, 50 µm-thick coronal brain sections were cut using a cryostat (RWD Minux FS800). For immunostaining experiments, tissue sections were rinsed in 1X PBS (3 times) and incubated in the blocking buffer (5 % Bovine Serum Albumin (BSA) + 0.5 % Triton X-100 + 1X PBS) (BSA-HIMEDIA Catalog No. TC194, Triton X-100 SRL Catalog No. 64518) for one hour at room temperature. Sections were then incubated in the primary antibody (dilution 1:1000 X in blocking buffer) at room temperature overnight. Sections were rinsed 3 times with 1X PBS + 0.5 % Triton X-100 solution and incubated for two hour in Alexa Fluor conjugated goat anti-rabbit/ chicken or donkey anti-goat/rabbit secondary antibodies (dilution 1:1000 X in blocking buffer) along with DAPI (SRL Catalog No.18668) at room temperature. Then sections were washed with 1X PBS + 0.5 % Triton X-100, and mounted onto charged glass slides (Ted Pella, Inc. Catalog No. 260382-3). Citifluor AF-1 mounting media (Ted Pella, Inc. Catalog No. 19470-1) was used to cover-slip (Blue star Microscope cover glass 24 x 60 mm 10 Gms) the slides. Subsequently, sections were imaged on the upright fluorescence microscope (Khush Enterprises, Bengaluru) (2X, 4X, and 10X lenses). ImageJ/FIJI processing software was used to process the images

### Dataset Creation and Augmentation

A curated behavioural video dataset was assembled to train and evaluate Scratcher, consisting of top-down recordings of mice engaged in acute, or spontaneous itch-related behaviours. Videos were captured using a custom-built videotaping chamber equipped with uniform backlighting to produce high-contrast silhouettes and recorded at 30 frames per second. A total of 17 behaviour videos were selected from independent experiments involving different itch models (chloroquine-induced acute itch and spontaneous behaviour). Scratches other than those directed towards the nape, such as those towards the cheeks, thorax, ears, etc., were labelled as “Others”. Subsequently, frames were extracted from these videos, and frames were annotated manually as “Locomotion”, “Others”, or “Itch” by two trained human annotators using Roboflow, an internet-browser-based labeling interface. Scratching was defined by rhythmic hindlimb movement when approaching or otherwise in contact with the nape, based on both speed and posture cues. Annotators labelling the videos were blind to the itch model in order to minimize and avoid any form of bias. To increase classifier robustness, we implemented the following augmentations to output 3 augmented images per training example - Flip (Horizontal, Vertical), 90° Rotate (Clockwise, Counter-Clockwise, Upside Down), Grayscale, Horizontal and Vertical flipping, Brightness (Between −15% and +15%) and Exposure (Between −10% and +10%). The full labeled dataset was then stratified and split into a training set (70%), a validation set (20%) and a test set (10%), while also ensuring that the class distribution of “Itch” versus Non-Itch (“Locomotion” and “Others”) was preserved in both subsets. The weights used in the initial phases of training were acquired from YOLOv8’s proprietary pre-trained models. Subsequent rounds of training were performed after giving due consideration to how Box_loss, Class_Loss and DFL_Loss evolved across epochs, depending on which, various parameters, including the training dataset, were shuffled and altered. Once the dataset was pre-processed and augmented, we exported the dataset in the JSON COCO segmentation format(Lin et al., 2014) with the training, validation, and test splits mentioned above. Additionally, given that the YOLOv8 architecture employs a “one-shot” detection algorithm, the dataset had to be curated such that it covers a diverse range of instances for each of the classes with as little redundancy as possible in order to avoid overfitting and overlearning. Therefore, to ensure that such instances are minimized, an early learning cutoff was set to prevent models from reaching asymptote training values.

### Heuristic for temporal Aggregation and Majority Classification implemented in Scratcher

To convert high-resolution frame-wise behavioural annotations into temporally aggregated second-wise summaries, a custom Python function *(filter_behaviours)* was implemented. This function processes frame-by-frame behavioural classifications obtained from the YOLOv8 detections model (recorded at 30 frames per second) and assigns a single behaviour label per second using a majority vote scheme, followed by heuristic corrections for composite behaviours. The input to the function is an *.xlsx* file containing frame-level behavioural annotations, with behaviours listed in the second column. A rolling bin of 30 consecutive rows (corresponding to one second of video at 30 fps) was used to segment the full dataset temporally. Within each one-second bin, a majority voting strategy determined the dominant behaviour. In cases where the most frequent behaviour was “Others” but the second most frequent was “Itch”, the behaviour was reassigned to “Itch” to account for the underrepresentation of transient itch events. Composite or co-occurring labels (e.g., “Itch, “Locomotion,” or “Locomotion, Others”) were clarified via empirically derived rule-based logic that favored biologically relevant prioritization (e.g., “Itch” was given precedence over “Others”). The aggregated behaviour labels were stored in a new DataFrame with columns “Seconds” (representing elapsed time in seconds from video start) and “Behaviour” (dominant behaviour during that second). This output was saved to a new Excel file with multiple sheets using *pandas* and *openpyxl*. A second sheet (“Behaviour Durations”) was generated to quantify the cumulative duration (in seconds) of each behaviour across the entire session. Behaviours not present in the session were explicitly added with a zero duration to ensure consistent output formatting across multiple files. The output included “Locomotion”, “Others”, “Itch”, and “no_detection” as predefined behaviour categories. To quantify itch-related activity over time, a binary vector (itch_counts) was created where each second was marked as 1 if “Itch” was the assigned label, and 0 otherwise. Rolling sum windows were applied to this binary vector to compute average itch occurrence per minute and per 3-minute intervals. These statistics were stored in a third sheet (“Itch Statistics”), which reported the average number of detected itch events per minute and per 3-minute bin across the full duration of the video.

### Gaussian Hidden-Markov Model Heuristic

This method leverages several Python libraries for data preprocessing, modeling, and evaluation. *Pandas* was used for structured data handling and merging the frame-wise predictions with second-wise ground truth. *NumPy* enabled efficient numerical operations and reshaping of label arrays. *Scikit-learn* (sklearn.preprocessing.LabelEncoder) was used to encode categorical behaviour labels into numerical form compatible with modeling tools. *hmmlearn* was employed to construct and train the Gaussian Hidden Markov Model, and *scikit-learn.metrics* provided functions such as accuracy_score and classification_report for model evaluation. We trained the Gaussian Hidden Markov Model (HMM) to infer the underlying behavioural state sequence for each 30-frame window. Each 30-frame behaviour sequence was encoded as integers using a fitted LabelEncoder. These were flattened and reshaped into a single column vector suitable for hmmlearn’s GaussianHMM, which models each observed behaviour as a Gaussian-distributed emission from a hidden state. The model was trained with the number of components set to the number of unique behaviours, using the lengths parameter to maintain the segmentation between 30-frame windows. For inference, we predicted the most frequent hidden state among the last five frames of the decoded sequence, providing a temporally smoothed and noise-robust prediction for the second-wise behaviour label. Per-frame behavioural predictions were first grouped into non-overlapping 1-second bins (30 frames per bin) based on frame indices. Ground-truth labels for each second were derived from manually annotated data. Bins were aligned such that only those with a corresponding ground truth label were used for training and evaluation. The full dataset contained 15,804 1-second bins, from which a subset was used in this experiment. After discarding frames from initial sections of the video to align with ground truth, 1-second bins were extracted from frame 81,000 onwards for predicted labels, and from second 2700 onwards for ground-truth annotations. Behaviours were string-labeled and mapped to integer-encoded states using a LabelEncoder. All unique behaviours from both the frame-wise predictions and ground-truth were used to fit the encoder. An unsupervised Gaussian HMM was trained using the hmmlearn library, with the following specifications: Number of hidden states - equal to the number of unique behaviours present, Covariance type - diagonal for assuming uncorrelated features, Observations - each 30-frame sequence was flattened and treated as a 1D time series of discrete behaviour encodings, Training input: all sequences were concatenated, and the lengths parameter was used to denote segment boundaries corresponding to each bin. The model was trained for 100 iterations with a fixed random seed for reproducibility (random_state=42). For each 30-frame input sequence, the trained HMM was used to infer the most probable state sequence using the Viterbi algorithm. To obtain a robust summary behaviour for the entire second, we employed a frequency-based heuristic: the most frequent state among the last 5 frames was selected as the representative behaviour for that 1-second bin. This tail-weighted heuristic biases the prediction toward the end of the second, reflecting our observation that scratch bouts often begin or end mid-window. Predicted state indices were then converted back into human-readable behaviour labels using the inverse transform of the fitted LabelEncoder. Accuracy was computed by comparing predicted 1-second behaviours against the ground-truth labels. We also computed a full classification report including precision, recall, and F1-score for each behaviour class. To ensure valid reporting, label-wise metrics were restricted to only those behaviours that were present in both the predictions and ground truth. A standalone utility function predict_behaviour() was implemented to allow prediction on novel 30-frame sequences. This function takes in raw per-frame behaviour labels, encodes them, and returns the predicted behaviour for the window based on the last inferred hidden state.

### Bayesian Model Heuristic

This method also employed a suite of Python libraries for data processing and statistical modeling. *Pandas* handled CSV loading and grouping of frame-wise behaviours into 30-frame windows. *NumPy* facilitated the manipulation of label arrays and ensured compatibility with evaluation metrics. *scikit-learn* (sklearn.preprocessing.LabelEncoder) was used to encode categorical labels into a numeric form. The Counter and defaultdict classes from Python’s built-in collections module were used to compute prior distributions and frame-wise behaviour counts. Model performance was assessed using accuracy_score and classification_report from *scikit-learn.metrics*. The Bayesian heuristic treated the prediction of a second-order behaviour as a classification task based on frame-wise counts in a 30-frame window. Prior probabilities for each behaviour were computed from the full dataset as the proportion of frames labeled with that behaviour. For each 30-frame window, we computed a pseudo-posterior probability for each candidate behaviour by multiplying the prior with a simple likelihood: if the behaviour was observed in the window, a proportional score based on its frequency (count / 30) was applied; if not, a small penalty factor (0.01) was used to reduce its likelihood. The behaviour with the highest resulting score was selected as the predicted label for that second. The resulting models were evaluated by comparing their second-wise predictions to the ground truth annotations. Accuracy and detailed class-wise metrics were computed using accuracy_score and classification_report from *scikit-learn*. To ensure fair comparison, only behaviour labels present in both predictions and ground truth were included in the evaluation.

### Aggregation heuristic using a sliding window, non-overlapping average probability

We implemented a heuristic method to assign second-wise behaviour labels based on confidence scores extracted from a YOLOv8-based behaviour detection pipeline. The process utilized a non-overlapping sliding window approach with a window size of 30 frames (corresponding to one second of video at 30 fps), ensuring that each behavioural prediction was derived from a distinct, non-overlapping chunk of time. The procedure was executed using various Python libraries. *Pandas* was used for CSV data manipulation, time binning, and group-level aggregation.NumPy (numpy)was used for array operations and imported wherever matrix-level operations were necessary. *Matplotlib.pyplot* and *Seaborn* were used for generating evaluation figures such as confusion matrices, metric bar plots, and scatter plots. *Scikit-learn* was utilized to compute evaluation metrics, including accuracy, precision, recall, and F1 score. The raw input generated from a YOLOv8 object detection model consisted of a CSV file with frame-wise behaviour predictions. Each row of the file included three fields: Frame (frame number), Behaviour (categorical label), and Confidence (YOLO-assigned score for detection reliability). For each second, all rows corresponding to the 30 frames in that second were grouped using group by(’Seconds’). Each second of video (corresponding to 30 frames at 30 frames per second) was treated as a discrete bin. Within each one-second window, we grouped the frames and calculated the average confidence score for each behaviour. The behaviour with the highest average confidence was selected as the representative label for that second. The behaviour with the highest average confidence was selected to represent that second. This rule maximized reliability by accounting for both frequency and confidence of detection within the window, thus increasing robustness to momentary misclassifications. The final result was a dataframe containing second-wise predictions, saved as a CSV file. The process was repeated for the entire video, producing a sequence of second-wise behaviour predictions. These were compared against ground truth annotations to evaluate performance.

### Aggregation heuristic using an overlapping sliding window

Frame-wise behavioural predictions were generated using YOLOv8, with each frame annotated with a behaviour label and corresponding confidence score. To convert these frame-level annotations into temporally smoothed predictions, we implemented a majority-vote-based sliding window approach using Python. This method aimed to reduce noise and transient misclassifications by leveraging the temporal continuity of behaviour. All analyses were performed in Python, utilizing the *pandas* library for data manipulation and the *scipy.stats* module to compute statistical modes. The raw output, comprising individual frame numbers and their corresponding predicted behaviour classes, was first preprocessed to ensure proper data types and sorted in order by frame index. Only frames with valid numerical indices were retained. A fixed length sliding window of 30 frames (equivalent to one second of video at 30 frames per second) was moved across the entire dataset with a stride of 1 frame. For each window, the mode (i.e., most frequently occurring behaviour) was computed. If a unique mode was found, it was assigned as the behaviour prediction for the window; in the rare event of a tie, the first occurring mode was selected. This behaviour label was then assigned to the frame at the center of the window, and the corresponding second was calculated based on its frame index. The process was repeated across the full length of the video data, generating a list of second-wise predictions smoothed by overlapping windowed voting. The resulting DataFrame, consisting of center-frame indices, their associated second, and predicted behaviours, was saved as a CSV file for downstream evaluation and comparison with ground truth annotations. These were compared against ground truth annotations to evaluate performance.

### Graphical User Interface (GUI)

A custom graphical user interface (GUI) was developed using Python’s *tkinter* library to streamline three sequential processes for rodent behaviour analysis from video data: (1) object detection using a YOLO-based model, (2) behaviour filtering from annotated frames, and (3) downstream behaviour analysis. The interface allows users to perform end-to-end processing, prediction, and training of YOLO models with minimal programming knowledge.*YOLO Model Integration.* Object detection was performed using the Ultralytics YOLO implementation. The GUI loads our custom-trained YOLO model using *ultralytics.YOLO(model_path)*. Video files (in .mp4 format) from a user-specified input directory are sequentially processed with this model. The function *process_video* - defined externally - handles frame-by-frame inference and result saving. Users can specify a confidence threshold (default 0.6), which determines the minimum detection probability for a prediction to be accepted. The results, typically exported as Excel spreadsheets containing frame-wise predictions, are saved to an output directory as specified by the user. Post-detection, output files from the inference step are processed by the function *filter_behaviours*, which applies user-defined rules to consolidate frame-wise labels into temporally contiguous behavioural bouts. The filtered outputs are saved as new Excel files prefixed with *raster_plot_input_*. The processed and filtered files are then passed to *analyse_behaviours*, a third function that performs downstream quantitative analyses such as bout duration calculations, transition probabilities, visualization of behavioural timelines, and a suite of other quantitative plots. This function is modular and can be tailored to the experimental context. *Prediction Mode.* For ad hoc predictions, a separate GUI tab enables inference on individual video files using a YOLO model. Users can optionally change the working directory before prediction. Output visualization is handled natively by YOLO.predict() method, with results saved if the “Save Output” checkbox is enabled. *Training Mode.* The interface also provides a tab for training custom YOLO models. The user specifies: A working directory where model weights and logs will be saved, A base model checkpoint (.pt format) to initialize training, A *data.yaml* file conforming to the Ultralytics YOLOv8 data format, which specifies training/validation paths and class names, and the number of training epochs. The training function uses *YOLO(model_path).train(data=data_path, epochs=n)* to initiate training. Time taken for training is logged to a text file *(Time_taken.txt)* in the working directory. All GUI components are built using *tkinter* and *ttk*, with structured layout management and modal dialogs for file selection, error handling, and user feedback. Error-checking routines are included to ensure path validity and input type correctness (e.g., float parsing for confidence, integer parsing for epochs). *Hardware Requirements.* CPU: Minimum 4-core processor (Intel i5/Ryzen 5 or higher recommended). RAM: Minimum 8 GB (16 GB recommended for large video processing or model training). GPU: NVIDIA GPU with CUDA support (Compute Capability ≥ 6.1) is highly recommended for YOLO inference and training. Models run significantly faster on a GPU compared to a CPU. *Software Requirements*. Operating System: Windows 10/11, macOS (with limitations on YOLO training), or any modern Linux distribution. Python Version: Python 3.8 or later. Python Package Dependencies: *ultralytics* (for YOLOv8): Install via pip install ultralytics, *tkinter*: Included by default with most Python distributions.pandas: For spreadsheet manipulation. Install via *pip install pandas*, *openpyxl*: Required for .xlsx I/O. Install via *pip install openpyxl, opencv-python*: Required for video processing if not handled inside *process_video*. Install via *pip install opencv-python.* Additional Files/Modules: *video_processing.py*: Must define the *process_video* function, accepting model, video path, output folder, video name, and confidence threshold.*behaviour_filtering.py*: Must define the filter_behaviours function. *behaviour_analysis.py*: Must define the analyse_behaviours function.

### Slope Plot

The dataset was stored in a Microsoft Excel spreadsheet, with the first column representing time (in minutes) and each subsequent column corresponding to scratch duration values for individual mice across multiple time points. All analyses and visualizations were conducted using Python, employing the following libraries: *pandas* (for data ingestion and handling), *numpy* (for numerical computations), *matplotlib.pyplot* (for plotting), and *scipy.stats.linregress* (for performing linear regression). The data were first imported using *pandas.read_excel*, assuming that the first row contained column headers. Time values from the first column were extracted as the independent variable, while each of the remaining columns was interpreted as the scratch behaviour profile of a single mouse. For each mouse, linear regression was performed using *scipy.stats.linregress*, modeling scratch duration as a linear function of time. The slope and intercept obtained from this regression were then used to compute predicted values across the observed time range. Only the regression lines were plotted, omitting raw scatter points to improve visual clarity and emphasize overall behavioural trends. The x-axis represented elapsed time in minutes and was fixed between 0 and 45 minutes, while the y-axis showed predicted scratch durations. Y-axis limits were dynamically scaled to exceed the maximum regression value by 10%, allowing for adequate separation of plotted lines.

### Peak Scratching Duration

To generate the scatter plot of peak scratching durations for individual mice, we used Python (version 3.10.12) along with the *pandas* (version 1.5.3), *matplotlib* (version 3.7.1), and *openpyxl* (for reading .xlsx files) libraries. The input was a Microsoft Excel spreadsheet with time-series data where the first column denoted time in minutes, and each subsequent column represented scratch duration data for an individual mouse. The dataset was loaded using *pandas.read_excel()* with the assumption that the first row of the file contained headers. The first column was parsed as the independent variable (time), while the rest of the columns were treated as dependent variables representing scratch durations for each mouse over time. For each mouse, the index corresponding to the maximum value in its respective column was computed using the *idxmax() method*, and both the peak scratching duration and the corresponding time point were extracted. These were stored as x- and y-coordinates for plotting, with mouse identifiers retained from the column headers for labeling. A scatter plot was constructed using *matplotlib.pyplot*, with each mouse’s peak duration represented as a single point, positioned according to the time since injection (x-axis) and peak scratch duration in seconds (y-axis). Distinct colors were assigned to each data point using the viridis colour map from *matplotlib.cm*, with the number of color bins matching the number of mice. The point size was fixed at 100 for visual clarity. The y-axis was scaled dynamically to accommodate 10% more than the maximum observed peak duration, while the x-axis was fixed from 0 to 45 minutes to reflect the duration of the behavioural assay.

### LOESS-Smoothened Time-Course Line Plots with SEM Shading

To examine the temporal dynamics of scratching behaviour across control and experimental groups when assaying for scratching behaviour, time-series data were processed and visualized using *Python* (version 3.10.12) with *pandas* (version 1.5.3), *numpy* (version 1.22.4), *matplotlib* (version 3.7.1), and scipy (version 1.10.1). Data were loaded from two Microsoft Excel files corresponding to baseline (control) and DCZ-treated (experimental) conditions. Each file contained columns for time (minutes), mean scratching duration, standard error of the mean (SEM), and the number of observations per time point. After loading the data using *pandas.read_excel(),* vectors corresponding to time, mean, and SEM were extracted for each group. To reduce noise and enhance the interpretability of behavioural trends, a cubic spline-based smoothing function was applied. Specifically, *scipy.interpolate.make_interp_spline()* was used to fit a third-degree spline to the data, and *numpy.linspace()* was used to interpolate 300 evenly spaced time points within the original range, yielding high-resolution smoothed curves. Smoothed mean scratching trajectories were plotted using *matplotlib.pyplot.plot()* with the control and experimental curves displayed in different colours. Shaded regions representing SEM around the original unsmoothed means were rendered using *plt.fill_between()* to visualize variability over time. Axes were labeled to indicate minutes post-injection and scratching duration in seconds. The figure was configured with a size of 8×5 inches, and a legend was included to distinguish between conditions. Plot styling followed the default matplotlib configuration, and grid lines were disabled for maximum visual clarity.

### Linear Regression and Correlation Analysis

To evaluate the agreement between human-annotated and model-annotated scratching durations, Pearson’s correlation coefficient was computed and visualized using a scatter plot. Data were imported from an Excel spreadsheet where the first and second columns represented the human and model annotations on the x and y axes, respectively. The Pearson correlation coefficient (LJ) was calculated to quantify linear agreement between the two sets of annotations. A scatter plot was generated in which each point represented a paired human-model measurement, and an identity line (slope = 1, intercept = 0) was overlaid to visualize perfect agreement. The Pearson LJ value was displayed within the plot area, and a minimalistic style was maintained with semi-transparent data points, gridlines, and appropriate axis labeling to facilitate interpretability.

### Heat-Map plot

To visualize the time-resolved changes in calcium activity, or scratching duration, across subjects, ΔF/F or scratching duration values were plotted as heatmaps using the Seaborn data visualization library in Python. Fluorescence data was imported from an Excel file, where the first column represented time points (in seconds/minutes) and each subsequent column contained the ΔF/F trace or the scratching duration for one subject (e.g., mouse). The data matrix was transposed such that rows represented individual subjects and columns corresponded to time points. This format enabled visual comparison of fluorescence/scratching dynamics across subjects over time. Heatmaps were generated using the *seaborn.heatmap()* function with a variety of diverging color palettes (’*coolwarm’*, ‘*seismic’*, ‘*RdBu_r’*, ‘*BrBG’*, ‘*PiYG’*, and ‘*Reds’*) to emphasize both increases and decreases in fluorescence or scratching durations. Depending on the user’s needs, colour maps that are sensitive to colour blind people have also been included. The diverging point of the colormap was explicitly set to zero to ensure symmetric visualization around baseline activity. Each heatmap was plotted with a color bar labeled “ΔF/F” or “Scratching Duration” to indicate the normalized intensity of either fluorescence or scratching duration. The x-axis was labeled with time (in seconds/minutes), and the y-axis denoted individual subjects. The *matplotlib.pyplot* library was used to configure the figure size and layout.

### Bland-Altman Analysis of Agreement

To assess the agreement between human-annotated and model-annotated scratching durations, a Bland-Altman analysis was performed. Scratch duration values were extracted from a spreadsheet containing paired annotations, with the second and third columns representing human and model values, respectively. The data were imported into Python using the *pandas* library. For each pair of values, the average was calculated as the mean of the human and model annotations, and the difference was calculated by subtracting the human annotation from the model prediction. These average and difference values formed the basis of the Bland-Altman plot, where each point represented a paired observation. The mean difference was computed to evaluate systematic bias between the two methods. Additionally, the standard deviation of the differences was calculated using a sample standard deviation (ddof=1). Limits of agreement were established at ±1.96 times the standard deviation around the mean difference, representing the 95% confidence interval assuming a normal distribution of differences. A scatter plot was then generated using *matplotlib*, with the average of the two methods on the x-axis and the difference (model minus human) on the y-axis. The plot included horizontal dashed lines representing the mean difference (bias) and the upper and lower limits of agreement. Points were displayed with semi-transparent teal colouring and outlined in black to ensure visibility. Grid lines were lightly overlaid to improve visibility

### False Positive rate and False Negative Rate

To assess the class-wise error characteristics of the model, false negative rates (FNR) and false positive rates (FPR) were computed individually for each behavioural class using Python 3.10 and the libraries *pandas*, *matplotlib*, and *seaborn*. This analysis was carried out using the merged frame-wise prediction dataframe *(comparison_df)*, which contains the ground truth labels *(Behaviour_truth)* and model-predicted labels *(Behaviour_test)* for each time point. For each unique behaviour present in the ground truth, the False Negative Rate (FNR) was calculated by first identifying all instances where the behaviour was correctly annotated in the ground truth. The number of false negatives was then defined as the count of time points where the ground truth label matched the behaviour of interest but the model-predicted label did not. FNR for that class was calculated as the ratio of false negatives to the total number of ground truth instances for that class:

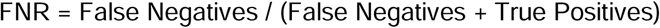

In parallel, the False Positive Rate (FPR) for each behaviour was computed by counting all instances where the behaviour was not present in the ground truth (i.e., all negative examples), and among them, how many times the model falsely labeled that behaviour. The FPR was thus calculated as:

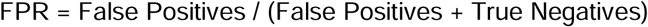

Both FNR and FPR values were aggregated into dictionaries indexed by behaviour class. These were subsequently visualized as bar plots using seaborn.barplot(). The FNR bar plot used the “Blues_d” colormap, while the FPR plot used “Reds_d”, visually separating the two error modalities. All plots were rendered using matplotlib.pyplot with a standard figure size of 10×5 inches, x-axis labels rotated for readability, and layout adjustments applied via plt.tight_layout().

### Area under the curve

To quantify cumulative scratching behaviour over time, the area under the curve (AUC) was calculated for each subject using Simpson’s rule for numerical integration. Scratch duration data were collected at regular time intervals for multiple animals and organized into a Microsoft Excel spreadsheet. The first column represented time (in minutes), while subsequent columns contained time-series scratch data for individual mice. The dataset was imported into Python using the *pandas* library. Time values were extracted as the shared x-axis, and each mouse’s scratch duration data was treated as an independent signal. Simpson’s rule, implemented via the *simps* function from the *scipy.integrate* module, was used to compute the AUC of each signal. Simpson’s rule was chosen over simpler numerical integration methods, such as the trapezoidal rule, because it fits a second-order (parabolic) polynomial to every pair of adjacent intervals. This makes it especially suitable for biological data that tends to exhibit curvature or non-linear transitions, as it captures the underlying dynamics more accurately than linear approximations. Compared to the trapezoidal method, Simpson’s rule reduces integration error when the signal contains smooth, continuous fluctuations, which is typical of time-resolved behavioural measures like scratching. The computed AUC values represent the total scratching activity integrated across the entire observation window for each animal. These values were visualized using bar plots generated with the *matplotlib* library. Each bar corresponded to an individual subject, the y-axis denoted the AUC, and the x-axis labeled each mouse. To ensure visual clarity, the y-axis was scaled to 10% of the maximum AUC, and x-axis labels were rotated to prevent overlap where necessary.

### Temporal Accuracy Analysis of Predicted behaviours

To quantitatively evaluate the performance of a behavioural classification algorithm, second-by-second predicted behaviour labels were compared against manually annotated ground truth labels over a 2700-second recording period. The analysis was implemented using *Python (v3.10)* with the *pandas*, *matplotlib*, and *seaborn* libraries. Two comma-separated value (CSV) files were loaded: one containing manually annotated behaviours (ground truth) and the other containing algorithmically predicted behaviours (test file). Both files were expected to include a time-aligned “Seconds” column and a categorical behaviour label column. The test file was truncated to the first 2700 rows to match the relevant temporal segment of the ground truth. Due to potential inconsistencies in timestamp indexing, the test file’s “Seconds” column was programmatically overwritten to ensure a sequential range from 1 to 2700, corresponding to one label per second. The ground truth and test datasets were then merged using an inner join on the “Seconds” column to align behaviour labels on a per-second basis. To avoid ambiguity from automatic merging suffixes, the resulting columns were explicitly renamed: the ground truth label was stored in *Behaviour_truth* and the predicted label in *Behaviour_test*. A new Boolean column, correct, was computed to denote whether the predicted label matched the ground truth at each time point. The overall classification accuracy was calculated as the mean of the correct column, representing the proportion of time points at which the prediction matched the annotated behaviour. Additionally, behaviour-wise accuracies were computed by grouping the merged dataframe by the ground truth label and calculating the mean correctness within each group. This allowed for the identification of differential model performance across behaviour categories. To visualize prediction reliability over time, a scatter plot was generated using *matplotlib*, where the x-axis represented time in seconds and the y-axis indicated whether each prediction was correct (1) or incorrect (0). This temporal representation helped identify regions of sustained accuracy or systematic misclassification. The plot was saved at 300 DPI resolution using lossless PNG format for high-quality reproduction in reports or publications.

### Confusion Matrix Analysis

To evaluate the categorical performance of the behaviour prediction model, a confusion matrix was constructed by comparing the predicted behaviour labels to ground truth annotations. The predicted labels were obtained from a CSV file containing model-inferred behaviours at each second (*Predicted_Behaviour*), while the ground truth labels were obtained from a corresponding CSV file annotated manually (*Behaviour*). The confusion matrix was computed using the confusion_matrix function from the *scikit-learn* (sklearn.metrics) library. Prior to comparison, both *dataframes* were temporally aligned using the Seconds column as a key, ensuring one-to-one correspondence between ground truth and predicted labels. The labels for the confusion matrix were explicitly defined using the unique set of behaviours present in the ground truth annotation to ensure consistent axis ordering and completeness in representation. The matrix was then visualized using a heatmap generated with the seaborn library (*sns.heatmap*). This heat map displayed the count of occurrences for each true vs. predicted label pair, with annotations enabled (annot=True) and formatted as integers (fmt=’d’). Color encoding was applied using the Blues colormap to enhance visual interpretation, and axis labels were set to reflect actual (y-axis) and predicted (x-axis) behaviour categories. The figure was saved at high resolution (dpi=300) in PNG format to a designated directory (save_dir), with tight bounding box settings to avoid truncation of labels or titles.

### Visualization of Class Distribution

To assess the class distribution and detect potential imbalances between annotated (ground truth) and model-predicted behavioural categories, horizontal count plots were generated for each dataset. This visualization aids in qualitative evaluation of class frequency agreement between manual annotations and automated predictions. Two CSV files were used as input: one containing the ground truth behaviour annotations *(ground_truth)* with a column labeled ‘Behaviour’, and the other containing the model-predicted behaviours *(test_file)* also under a ‘Behaviour’ or equivalent column. Both data frames were assumed to be temporally aligned at the per-second resolution. The class order for both plots was standardized by using the frequency-sorted order of behaviours from the ground truth dataset. This was accomplished by passing *order=ground_truth[’Behaviour’].value_counts().index* to the *sns.countplot()* function, ensuring consistent axis alignment and facilitating direct visual comparison. Visualization was performed using *seaborn* in conjunction with *matplotlib*. A side-by-side subplot layout *(plt.subplot(1, 2, i))* was used to juxtapose the ground truth distribution against the model prediction distribution. Both plots were set to horizontal orientation (y=’Behaviour’) to maximize readability of categorical labels, especially when label names are long or numerous. The combined figure was rendered at a width of 12 inches and height of 6 inches (figsize=(12, 6)), and exported at 300 DPI resolution in PNG format to a specified directory (save_dir) using *plt.savefig().* A tight layout adjustment *(plt.tight_layout())* was applied to prevent overlapping of elements.

### Classifier Performance Evaluation and Visualization

To quantitatively assess the performance of the behavioural classification algorithm, standard supervised classification metrics—accuracy, precision, recall, and F1-score—were computed using the merged ground truth and prediction dataset (comparison_df). These metrics provide a comprehensive overview of both the overall and class-specific predictive quality of the model. All calculations were conducted using functions from the sklearn.metrics module (Scikit-learn version ≥0.24 recommended). The accuracy_score() function was used to determine the proportion of correctly predicted labels across all classes. Weighted versions of precision_score(), recall_score(), and f1_score() were employed using average=“weighted” to account for class imbalance by weighting each class’s contribution by its support (i.e., number of true instances). The ground truth labels (Behaviour_truth) and predicted labels (Behaviour_test) were compared at each time point (typically seconds), and correctness was stored as a boolean column (correct) in the merged DataFrame. The values of all four metrics were rounded to two decimal places and printed for inspection. To enhance interpretability and facilitate presentation, the computed metrics were visualized using an interactive bar plot generated via plotly.express (Plotly version ≥5.0). A separate DataFrame (metrics_df) was created with two columns: “Metric” and “Value”, listing the four performance indicators. The px.bar() function was used to render the plot with distinct color encoding (color=“Metric”) and the Vivid color scheme from px.colors.qualitative to ensure high contrast and clarity. Metric values were annotated directly above each bar using *textposition=“outside“* and displayed as percentages by formatting the y-axis with *yaxis_tickformat=”.0%”.* The resulting figure was exported as a high-resolution PNG image using *fig.write_image()* and saved to the user-specified output directory *(save_dir)*.

### Peri-event aligned extraction of fluorescence data from annotated scratching events

This was done using custom Python scripts (Python ≥ 3.8, pandas v1.5+, numpy v1.22+). Fluorescence signals were acquired from CSV files containing timestamped CH1 (GCaMP) Z-score values, recorded via fiber photometry. The timestamps, originally in milliseconds, were converted to seconds to match the temporal resolution of the behavioural annotations generated by our automated labeling tool, Scratcher. These annotations consisted of second-level behavioural labels, including the “Itch” class. We extracted all timepoints labeled as “Itch” and performed a peri-event windowing procedure around each instance. To prevent over-representation of closely spaced itch events, we defined bouts of scratching by grouping consecutive “Itch” events occurring within 3 seconds of each other. If a gap of more than 3 seconds occurred between two events, a new bout was initiated.The first scratch in each (sub-)bout was used as the alignment point for peri-event analysis. For every itch event, we defined a symmetric window spanning from 5 seconds before to 5 seconds after the onset of scratching (−5s to +5s relative to event time). Since the fiber photometry system recorded at approximately 30 Hz, each second-long bin was expected to contain up to 30 data points. To ensure consistency, all fluorescence timepoints were rounded to six decimal places. For each second in the 10-second window, CH1 values were grouped by their integer second value. If more than 30 samples were available for a given second, only the first 30 were retained. If fewer than 30 samples were present, the row was padded with *NaN* values to maintain consistent dimensionality. These values were stored in row-wise dictionaries, mapping each aligned second to its vector of fluorescence values. The resulting output was stored as a tabular file, with each row representing a peri-event window aligned to an individual itch event.

### Raster Plot

To visualize the temporal distribution of discrete behavioural events across multiple subjects, a behavioural raster plot was generated using custom Python scripts. This method allowed for the simultaneous representation of distinct behaviours for multiple animals over a shared time axis, aiding in qualitative inspection of behavioural patterns, co-occurrence, and transitions. A behavioural log containing timestamped event information was prepared in Microsoft Excel format. The first column contained timepoints (in seconds), and each subsequent column represented the binary state (1 for event occurrence, 0 for absence) of a specific behaviour in an individual mouse. The dataset was imported using the pandas library, and temporal boundaries were identified by computing the minimum and maximum time values from the “Time” column. The behavioural raster was created using the matplotlib plotting library, with each subject represented by a horizontal row stacked vertically. The broken_barh function was employed to render behaviour-specific segments as horizontal bars aligned with their corresponding time intervals. For each subject, a list of (start time, duration) tuples was constructed from the raw data, where event durations were inferred from consecutive non-zero entries in the time series. Each behaviour was color-coded using a predefined colormap, ensuring visual distinction between behavioural states. The vertical height of each raster row was controlled using a row_height parameter, and the behaviours were labeled on the y-axis using plt.yticks, corresponding to the stacking order. The x-axis denoted the shared temporal domain (in seconds), with ticks placed at regular intervals. The plot was rendered with a dark background using the plt.style.use(’dark_background’) setting to enhance visual contrast. All other font elements, including tick labels and axis titles, were bolded for legibility.

### Statistical analysis

All statistical analyses were performed using GraphPad PRISM 8.0.2 software. Student t-tests were performed wherever applicable. ns > 0.05, ∗ P ≤ 0.05, ∗∗ P ≤ 0.01, ∗∗∗ P ≤ 0.001, ∗∗∗∗ P ≤ 0.0005.

## Illustration drawing

Cartoons with mice were made partially in BioRender (BioRender, 2022).

## Data Availability

The codes for installing the GUI and generating the analyses can be downloaded from GitHub https://github.com/BarikLab-IISc/Scratcher

## Acknowledgements

We thank the Barik lab members for their help and support. We would also like to thank Mr. Annappaswamy for his continued help with animal colony management. This work was supported by IndiaAlliance Intermediate Fellowship awarded to A.B. and IISc.

---

**Supplementary Figure 1.**
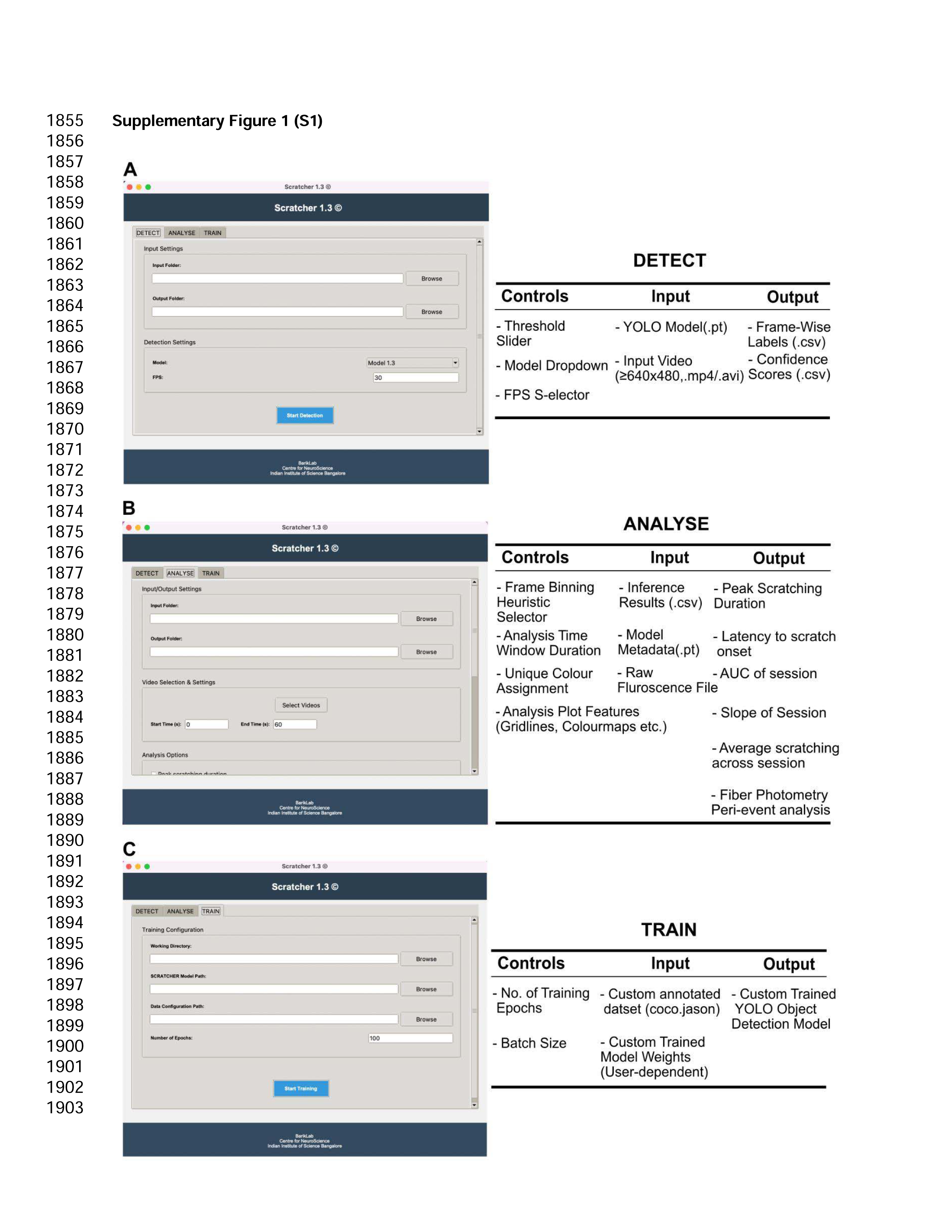
Graphical overview of the input-control-output architecture across the DETECT, ANALYSE, and TRAIN modules of the automated scratching detection and behavioral analysis tool. (A) The DETECT module allows users to input a video file and YOLO-based model weights to perform frame-wise behavioral classification, with adjustable parameters such as confidence thresholds, model selection, and frame rate settings. The output includes label and confidence score files in CSV format. (B) The ANALYSE module enables parameterized behavioral analysis using model outputs and raw fluorescence data. Controls include heuristic selection, analysis window configuration, and plotting options. Outputs include summary behavioral metrics such as scratching duration, onset latency, session slope, and photometry-based peri-event measures. (C) The TRAIN module provides customization for training object detection models using user-annotated datasets. Inputs include COCO-formatted datasets and custom training parameters. Outputs are user-trained YOLO models and associated weights.

**Supplementary Figure 2.**
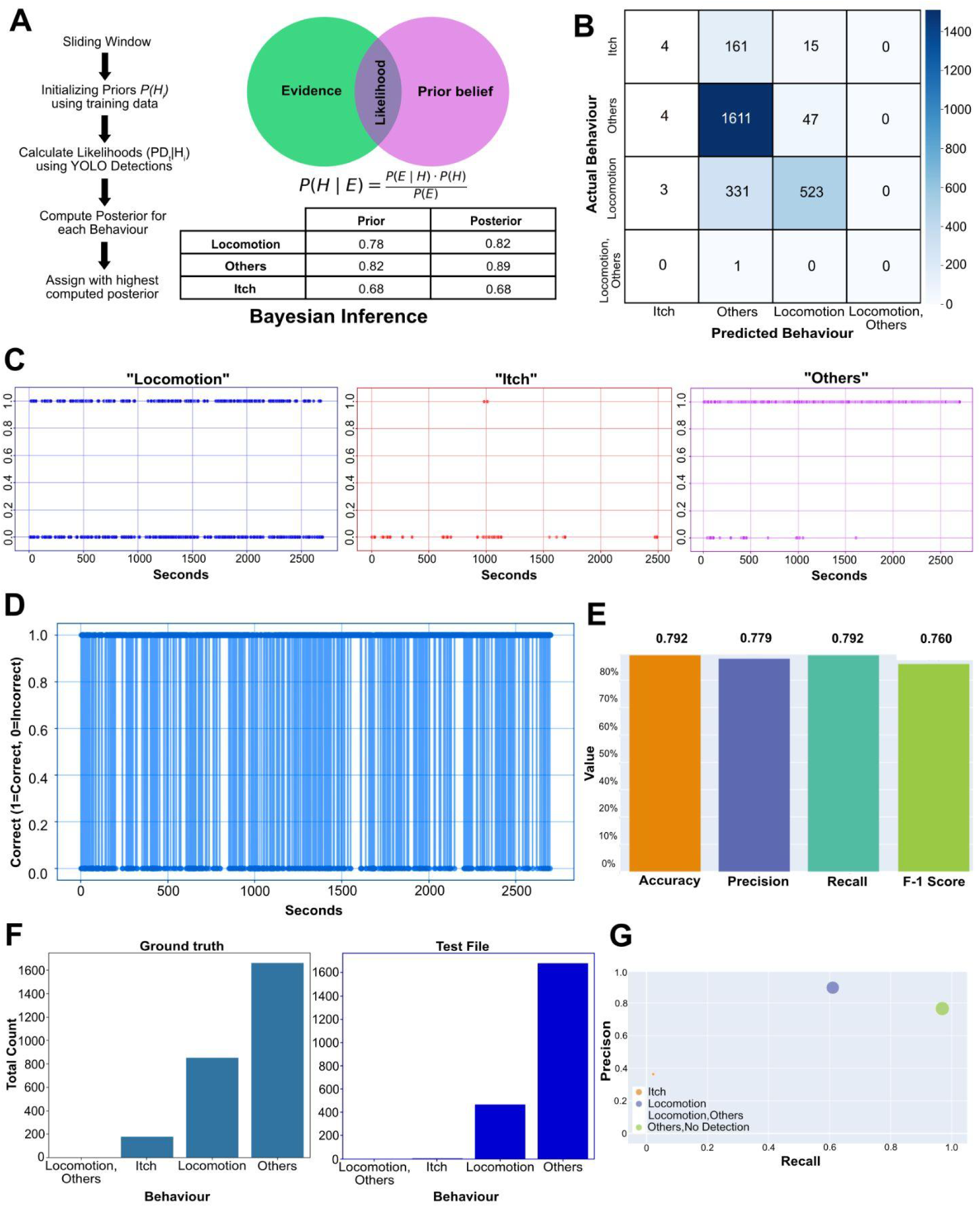
Performance evaluation of the Bayesian Inference heuristic for post-hoc binning for Frame-to-Second behavioural aggregation using YOLOv8 frame-wise outputs. (A) Bayesian inference-based aggregation schematic. A diagram illustrating the Bayesian approach to second-wise classification. Each sliding window aggregates YOLO detections to compute likelihoods, which are then combined with empirical priors to calculate posteriors. The behaviour with the highest posterior is assigned to the window. A representative table shows sample priors and posteriors for each class. (B) Confusion matrix comparing ground truth vs. test file labels. Class-wise distribution of predictions from the Bayesian method against ground truth annotations. (C) Accuracy of temporal distribution of individual classes. Scatter plot showing the correctness of “Locomotion”, “Itch” and “Others” (left to right) behaviour predictions over time, with y = 1 indicating agreement with ground truth (correct) and y = 0 indicating disagreement (incorrect). Each point represents a time-stamped prediction. (D) Temporal accuracy over the session. Each blue dot represents a second where the predicted label matches the ground truth (1 = correct, 0 = incorrect). The pattern reveals moments of behavioural misclassification and overall temporal fidelity. (E) Aggregate classification metrics. Accuracy, precision, recall, and F1-score are shown. (F) behavioural distribution comparison. Bar plots showing the total count of each behaviour type in the ground truth and test file. (G) Per-class precision vs. recall. Scatter plot showing precision and recall for each behavioural class.

**Supplementary Figure 3.**
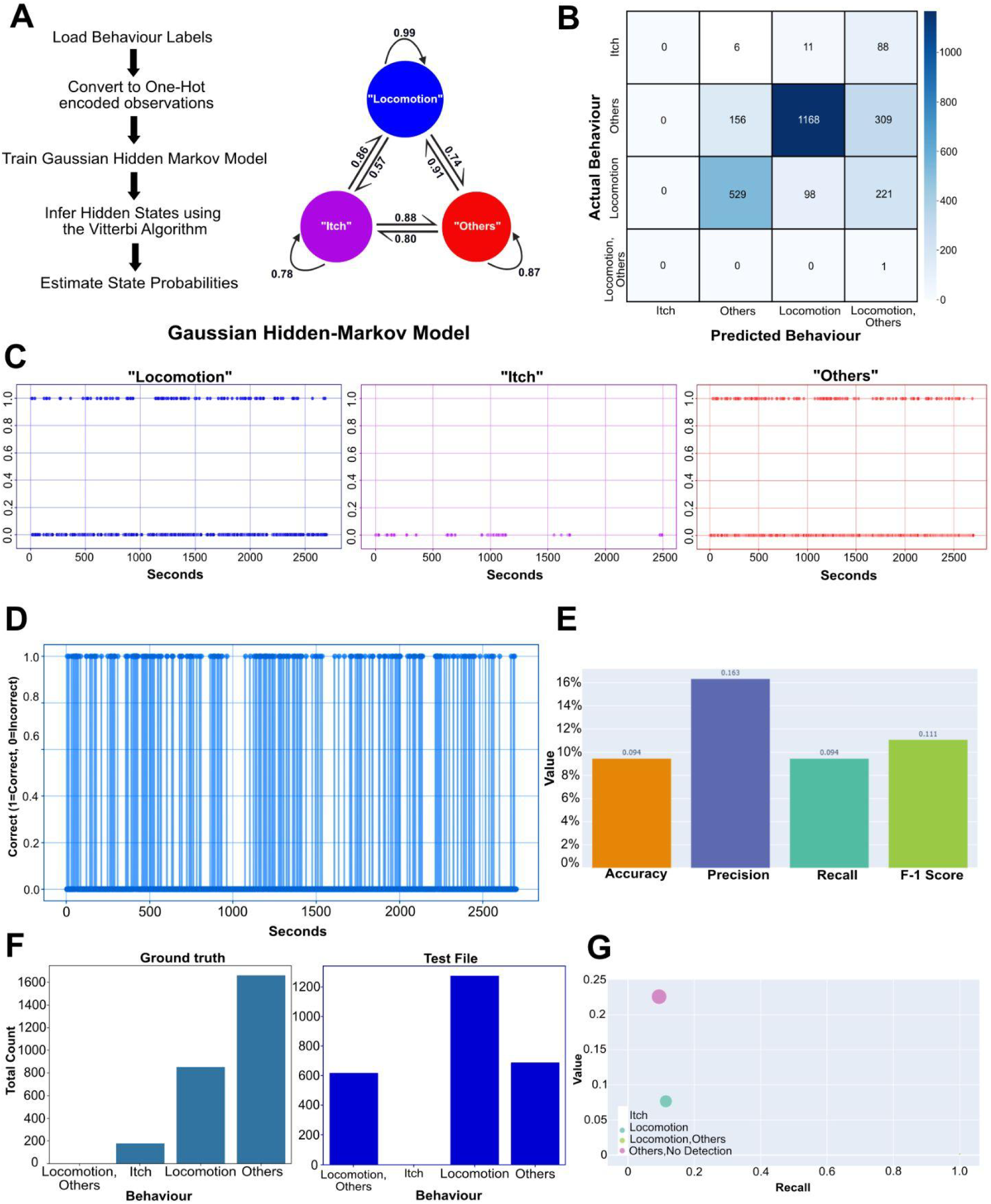
Performance evaluation of the Gaussian Hidden Markov Model (HMM) heuristic for post-hoc binning for Frame-to-Second behavioural aggregation using YOLOv8 frame-wise outputs. (A) Workflow and state transition diagram for HMM-based inference. A schematic describing the HMM-based approach. Frame-wise YOLOv8 predictions are converted to one-hot encoded vectors, used to train a Gaussian HMM. The Viterbi algorithm infers the sequence of hidden states, estimating the most probable behavioural labels. The right sub-panel shows the learned state transition probabilities between “Locomotion,” “Itch,” and “Others.“ (B) Confusion matrix comparing ground truth vs. HMM-based predictions. (C) Accuracy of the temporal distribution of individual classes. Scatter plot showing the correctness of “Locomotion”, “Itch” and “Others” (left to right) behaviour predictions over time, with y = 1 indicating agreement with ground truth (correct) and y = 0 indicating disagreement (incorrect). Each point represents a time-stamped prediction. (D) Temporal classification accuracy across the session. Binary sequence indicating per-second correctness (1 = correct, 0 = incorrect). (E) Aggregate classification metrics. Accuracy, precision, recall, and F1-score are shown. (F) Behavioural distribution comparison. Bar plots comparing the total number of each behaviour in the ground truth (left) and HMM output (right). (G) Per-class performance comparison. Scatter plot of precision vs. recall for each behavioural class.

**Supplementary Figure 4.**
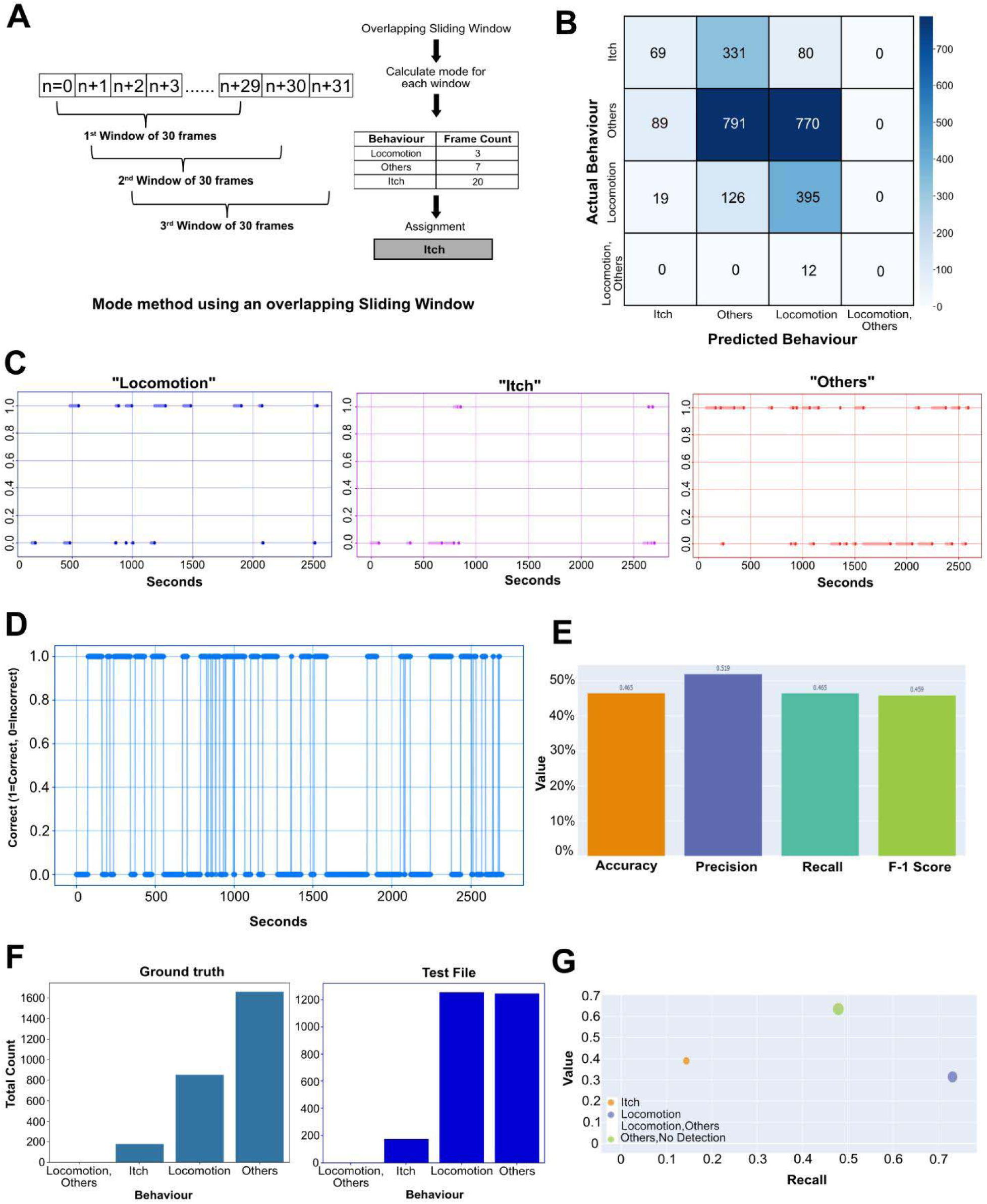
Performance evaluation of the overlapping sliding window heuristic for post-hoc binning to achieve Frame-to-Second behavioural aggregation using YOLOv8 frame-wise outputs. (A) Workflow of the overlapping sliding window heuristic. A 30-frame overlapping window moves across YOLOv8 frame-wise outputs (n to n+29). For each window, the mode of the behaviour classes is computed and assigned as the second-wise label. The accompanying table shows an example of frame counts per class and how “Itch” is chosen by majority. (B) Confusion matrix comparing ground truth vs. heuristic predictions. This panel shows the number of true vs. predicted behaviour labels. (C) Accuracy of the temporal distribution of individual classes. Scatter plot showing the correctness of “Locomotion”, “Itch”, and “Others” (left to right) behaviour predictions over time, with y = 1 indicating agreement with ground truth (correct) and y = 0 indicating disagreement (incorrect). Each point represents a time-stamped prediction. (D) Temporal classification accuracy across the session. Binary sequence indicating per-second correctness (1 = correct, 0 = incorrect). (E) Aggregate classification metrics. Accuracy, precision, recall, and F1-score are shown. (F) Comparison of behavioural distribution. The left panel shows the ground truth behaviour counts, while the right panel shows the distribution produced by the sliding window method. (G) A per-class scatter plot of precision vs. recall.

**Supplementary Figure 5.**
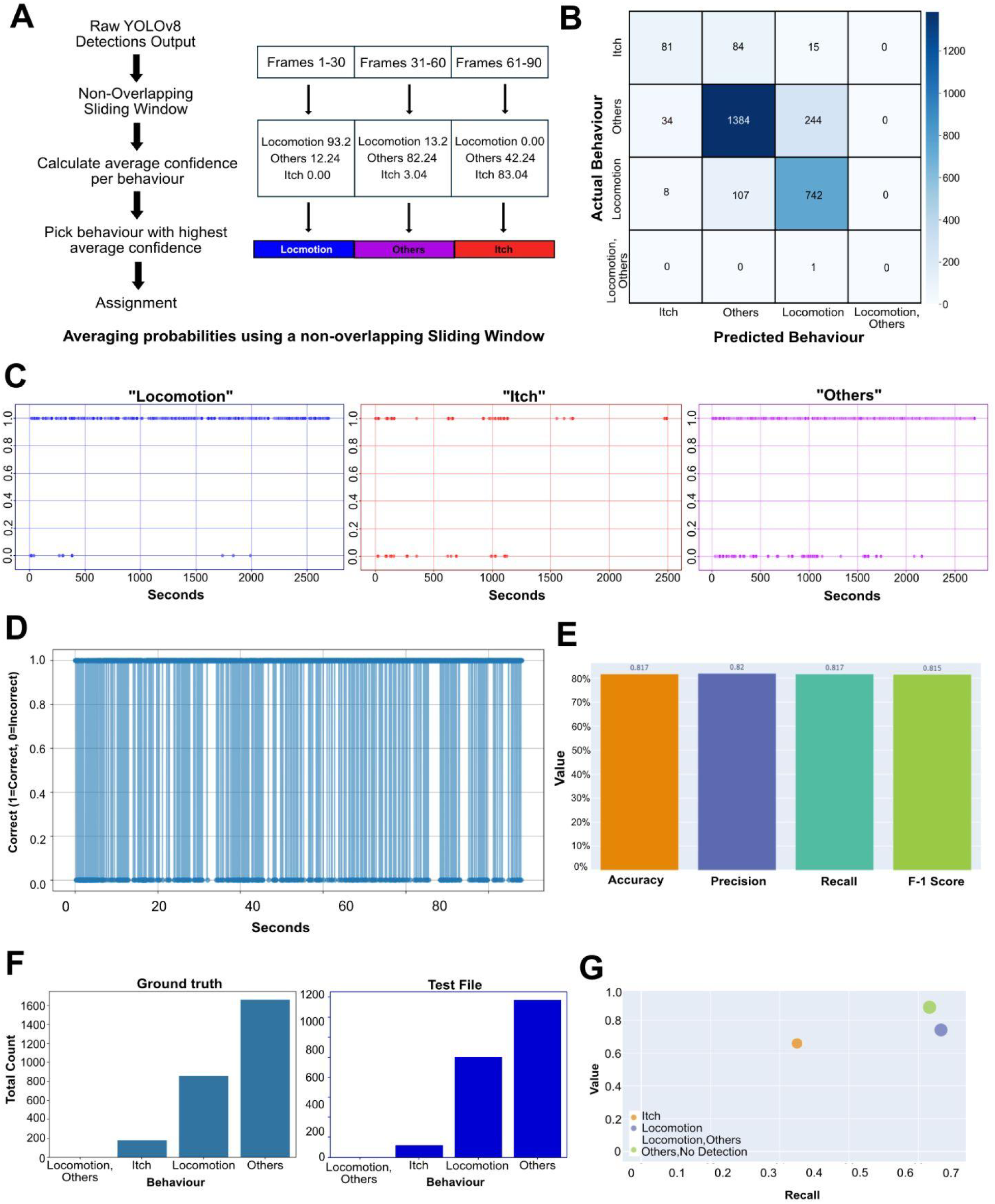
Performance evaluation of a non-overlapping sliding window-based probability averaging heuristic for post-hoc binning to achieve Frame-to-Second behavioural aggregation using YOLOv8 frame-wise outputs. (A) Schematic of the non-overlapping sliding window-based probability averaging heuristic. YOLOv8 frame-wise confidence values for each behaviour class are grouped into non-overlapping windows of 30 frames. For each window, average confidence scores are computed for all classes, and the class with the highest average is selected. An example shows how three consecutive windows yield predictions of “Locomotion,” “Others,” and “Itch.“ (B) Confusion matrix comparing predicted vs. true behaviours. (C) Accuracy of the temporal distribution of individual classes. Scatter plot showing the correctness of “Locomotion”, “Itch”, and “Others” (left to right) behaviour predictions over time, with y = 1 indicating agreement with ground truth (correct) and y = 0 indicating disagreement (incorrect). Each point represents a time-stamped prediction. (D) Second-wise correctness trace over time. A plot marking correct (1) or incorrect (0) predictions per second across a 2500-second test segment. (E) Aggregate classification metrics. (F) Comparison of behaviour distributions. The left panel shows ground truth counts for each behaviour; the right panel shows predictions using this heuristic. (G) Class-wise precision vs. recall scatter plot. Each dot represents a behaviour class.

**Supplementary Figure 6.**
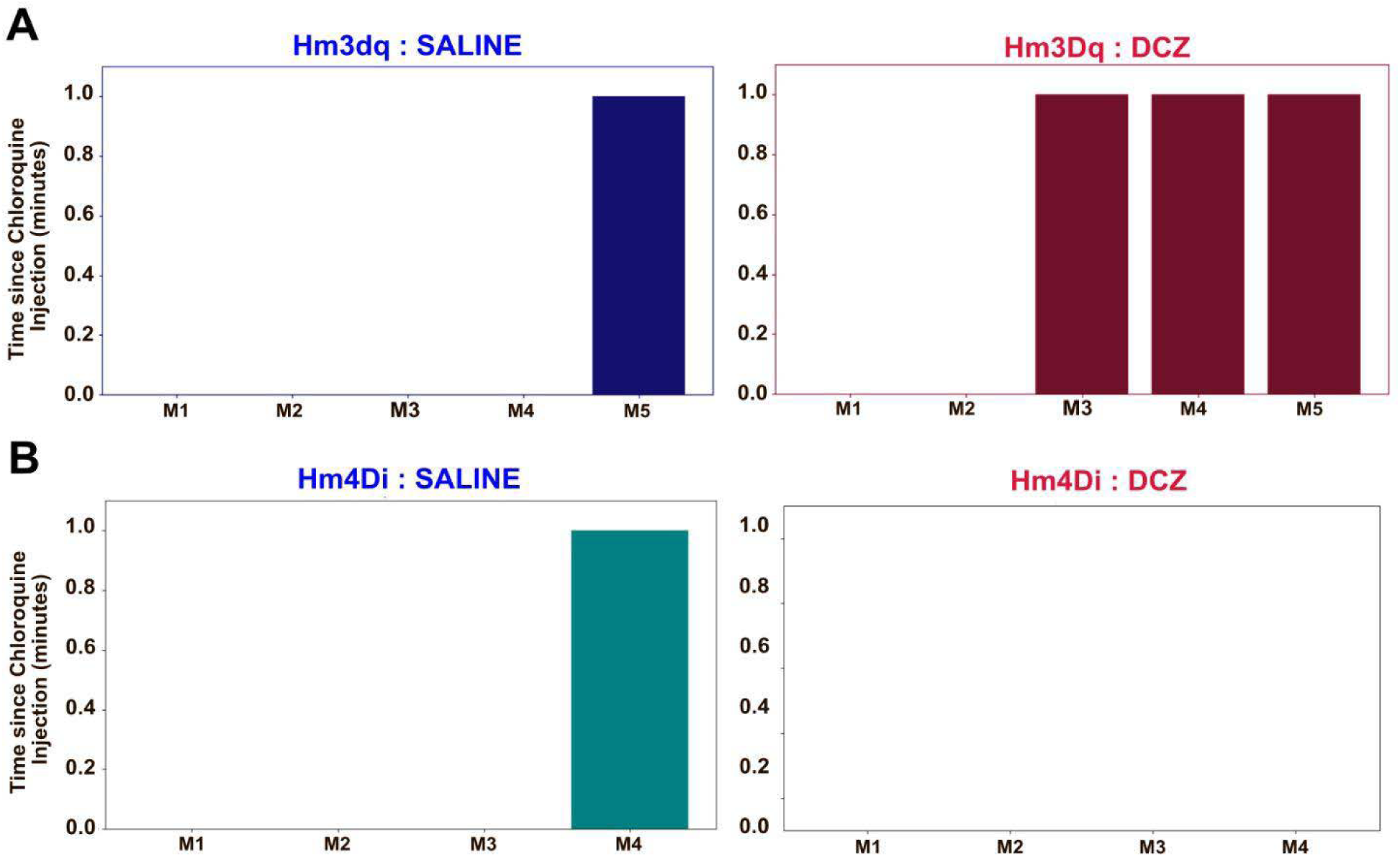
Latency to Itch Across Experimental Conditions. Plots showing the latency to first itch response following chloroquine injection across multiple experimental groups. (A) Comparison between Hm3Dq-expressing mice administered DCZ versus saline. (B) Comparison between Hm4Di-expressing mice administered DCZ versus saline.

## References

Argiento R, Lanzarone E, Villalobos IA, Mattei A, editors. 2017. Bayesian statistics in action, 1st ed, Springer proceedings in mathematics & statistics. Cham, Switzerland: Springer International Publishing. doi:10.1007/978-3-319-54084-9

Beyer HL, Morales JM, Murray D, Fortin M-J. 2013. The effectiveness of Bayesian state-space models for estimating behavioural states from movement paths. Methods Ecol Evol 4:433–441. doi:10.1111/2041-210x.12026

Brash HM, McQueen DS, Christie D, Bell JK, Bond SM, Rees JL. 2005. A repetitive movement detector used for automatic monitoring and quantification of scratching in mice. J Neurosci Methods 142:107–114. doi:10.1016/j.jneumeth.2004.08.001

Chen H, Feng R, Wu S, Xu H, Zhou F, Liu Z. 2022. 2D Human Pose Estimation: A Survey. *arXiv [csCV]*. Chen X-J, Sun Y-G. 2020. Central circuit mechanisms of itch. Nat Commun 11:3052. doi:10.1038/s41467-020-16859-5

Chun KS, Kang YJ, Lee JY, Nguyen M, Lee B, Lee R, Jo HH, Allen E, Chen H, Kim J, Yu L, Ni X, Lee K, Jeong H, Lee J, Park Y, Chung HU, Li AW, Lio PA, Yang AF, Fishbein AB, Paller AS, Rogers JA, Xu S. 2021. A skin-conformable wireless sensor to objectively quantify symptoms of pruritus. Sci Adv 7. doi:10.1126/sciadv.abf9405

Dang M, Liu G, Xu Q, Li K, Wang D, He L. 2024. Multi-object behavior recognition based on object detection for dense crowds. Expert Syst Appl 248:123397. doi:10.1016/j.eswa.2024.123397

DeNardo LA, Liu CD, Allen WE, Adams EL, Friedmann D, Fu L, Guenthner CJ, Tessier-Lavigne M, Luo L. 2019. Temporal evolution of cortical ensembles promoting remote memory retrieval. Nat Neurosci 22:460–469. doi:10.1038/s41593-018-0318-7

Deuis JR, Dvorakova LS, Vetter I. 2017. Methods used to evaluate pain behaviors in rodents. Front Mol Neurosci 10:284. doi:10.3389/fnmol.2017.00284

Doğan NÖ. 2018. Bland-Altman analysis: A paradigm to understand correlation and agreement. Turk J Emerg Med 18:139–141. doi:10.1016/j.tjem.2018.09.001

Elliott GR, Vanwersch RA, Bruijnzeel PL. 2000. An automated method for registering and quantifying scratching activity in mice: use for drug evaluation. J Pharmacol Toxicol Methods 44:453–459. doi:10.1016/s1056-8719(01)00111-3

Elliott P, G’Sell M, Snyder LM, Ross SE, Ventura V. 2017. Automated acoustic detection of mouse scratching. PLoS One 12:e0179662. doi:10.1371/journal.pone.0179662

Ge F, Xuan K, Lou P, Li J, Jiang L, Wang J, Lin Q. 2024. Multi-object detection and behavior tracking of sea cucumbers with skin ulceration syndrome based on deep learning. Front Mar Sci 11. doi:10.3389/fmars.2024.1365155

Guenthner CJ, Miyamichi K, Yang HH, Heller HC, Luo L. 2013. Permanent genetic access to transiently active neurons via TRAP: targeted recombination in active populations. Neuron 78:773–784. doi:10.1016/j.neuron.2013.03.025

Hussain M. 2024. YOLOv5, YOLOv8 and YOLOv10: The go-to detectors for real-time vision. arXiv [csCV].

Hu Y, Ferrario CR, Maitland AD, Ionides RB, Ghimire A, Watson B, Iwasaki K, White H, Xi Y, Zhou J, Ye B. 2023. : Quantification of user-defined animal behaviors using learning-based holistic assessment. Cell Rep Methods 3:100415. doi:10.1016/j.crmeth.2023.100415

Kielbinski M, Bernacka J. 2024. Fiber photometry in neuroscience research: principles, applications, and future directions. Pharmacol Rep 76:1242–1255. doi:10.1007/s43440-024-00646-w

Kobayashi K, Matsushita S, Shimizu N, Masuko S, Yamamoto M, Murata T. 2021. Automated detection of mouse scratching behaviour using convolutional recurrent neural network. Sci Rep 11:658. doi:10.1038/s41598-020-79965-w

Le Bras A. 2023. A new system to study mouse scratching behavior. Lab Anim (NY) 52:32. doi:10.1038/s41684-023-01114-3

Li J-N, Ren J-H, He C-B, Zhao W-J, Li H, Dong Y-L, Li Y-Q. 2021. Projections from the lateral parabrachial nucleus to the lateral and ventral lateral periaqueductal gray subregions mediate the itching sensation. Pain 162:1848–1863. doi:10.1097/j.pain.0000000000002193

Lin T-Y, Maire M, Belongie S, Hays J, Perona P, Ramanan D, Dollár P, Zitnick CL. 2014. Microsoft COCO: Common objects in contextComputer Vision – ECCV 2014, Lecture Notes in Computer Science. Cham: Springer International Publishing. pp. 740–755. doi:10.1007/978-3-319-10602-1_48

Ludbrook J. 2010. Confidence in Altman-Bland plots: a critical review of the method of differences. Clin Exp Pharmacol Physiol 37:143–149. doi:10.1111/j.1440-1681.2009.05288.x

Macdonald IL, Raubenheimer D. 1995. Hidden Markov models and animal behaviour. Biom J 37:701–712. doi:10.1002/bimj.4710370606

Mathis A, Mamidanna P, Cury KM, Abe T, Murthy VN, Mathis MW, Bethge M. 2018. DeepLabCut: markerless pose estimation of user-defined body parts with deep learning. Nat Neurosci 21:1281–1289. doi:10.1038/s41593-018-0209-y

McNamara JM, Green RF, Olsson O. 2006. Bayes’ theorem and its applications in animal behaviour. Oikos 112:243–251. doi:10.1111/j.0030-1299.2006.14228.x

Mu D, Deng J, Liu K-F, Wu Z-Y, Shi Y-F, Guo W-M, Mao Q-Q, Liu X-J, Li H, Sun Y-G. 2017. A central neural circuit for itch sensation. Science 357:695–699. doi:10.1126/science.aaf4918

Nagai Y, Miyakawa N, Takuwa H, Hori Y, Oyama K, Ji B, Takahashi M, Huang X-P, Slocum ST, DiBerto JF, Xiong Y, Urushihata T, Hirabayashi T, Fujimoto A, Mimura K, English JG, Liu J, Inoue K-I, Kumata K, Seki C, Ono M, Shimojo M, Zhang M-R, Tomita Y, Nakahara J, Suhara T, Takada M, Higuchi M, Jin J, Roth BL, Minamimoto T. 2020. Deschloroclozapine, a potent and selective chemogenetic actuator enables rapid neuronal and behavioral modulations in mice and monkeys. Nat Neurosci 23:1157–1167. doi:10.1038/s41593-020-0661-3

Oksuz K, Cam BC, Kalkan S, Akbas E. 2021. Imbalance problems in object detection: A review. IEEE Trans Pattern Anal Mach Intell 43:3388–3415. doi:10.1109/TPAMI.2020.2981890

Onishi T, Hirose K, Sakaba T. 2024. Molecular tools to capture active neural circuits. Front Neural Circuits 18:1449459. doi:10.3389/fncir.2024.1449459

Patterson TA, Basson M, Bravington MV, Gunn JS. 2009. Classifying movement behaviour in relation to environmental conditions using hidden Markov models. J Anim Ecol 78:1113–1123. doi:10.1111/j.1365-2656.2009.01583.x

Pavlenko D, Ishida H, Markan A, Akiyama T. 2025. A subpopulation of projections from the parabrachial nucleus to the central amygdala mediates itch. Sci Rep 15:26432. doi:10.1038/s41598-025-08612-z

Pereira TD, Tabris N, Matsliah A, Turner DM, Li J, Ravindranath S, Papadoyannis ES, Normand E, Deutsch DS, Wang ZY, McKenzie-Smith GC, Mitelut CC, Castro MD, D’Uva J, Kislin M, Sanes DH, Kocher SD, Wang SS-H, Falkner AL, Shaevitz JW, Murthy M. 2022. SLEAP: A deep learning system for multi-animal pose tracking. Nat Methods 19:486–495. doi:10.1038/s41592-022-01426-1

Piyush Shah D, Barik A. 2022. The spino-parabrachial pathway for itch. Front Neural Circuits 16:805831. doi:10.3389/fncir.2022.805831

Prajapati JN, Reddy P, Barik A. 2024. Neural pathways that compel us to scratch an itch. J Biosci 49. doi:10.1007/s12038-024-00452-9

Prajapati JN, Shah DP, Barik A. 2025. An intra-brainstem circuitry for pain-induced inhibition of itch. Neuroscience 568:95–107. doi:10.1016/j.neuroscience.2025.01.008

Redmon J, Divvala S, Girshick R, Farhadi A. 2016. You only look once: Unified, real-time object detection2016 IEEE Conference on Computer Vision and Pattern Recognition (CVPR). Presented at the 2016 IEEE Conference on Computer Vision and Pattern Recognition (CVPR). IEEE. doi:10.1109/cvpr.2016.91

Ren J, Shen X, Lin Z, Mech R. 2020. Best Frame Selection in a Short VideoProceedings of the IEEE/CVF Winter Conference on Applications of Computer Vision. pp. 3212–3221.

Ren X, Liu S, Virlogeux A, Kang SJ, Brusch J, Liu Y, Dymecki SM, Han S, Goulding M, Acton D. 2023. Identification of an essential spinoparabrachial pathway for mechanical itch. Neuron 111:1812–1829.e6. doi:10.1016/j.neuron.2023.03.013

Sakamoto N, Haraguchi T, Kobayashi K, Miyazaki Y, Murata T. 2022. Automated scratching detection system for black mouse using deep learning. Front Physiol 13:939281. doi:10.3389/fphys.2022.939281

Samineni VK, Grajales-Reyes JG, Grajales-Reyes GE, Tycksen E, Copits BA, Pedersen C, Ankudey ES, Sackey JN, Sewell SB, Bruchas MR, Gereau RW. 2021. Cellular, circuit and transcriptional framework for modulation of itch in the central amygdala. Elife 10. doi:10.7554/eLife.68130

Samineni VK, Grajales-Reyes JG, Sundaram SS, Yoo JJ, Gereau RW 4th. 2019. Cell type-specific modulation of sensory and affective components of itch in the periaqueductal gray. Nat Commun 10:4356. doi:10.1038/s41467-019-12316-0

Schweihoff JF, Loshakov M, Pavlova I, Kück L, Ewell LA, Schwarz MK. 2021. DeepLabStream enables closed-loop behavioral experiments using deep learning-based markerless, real-time posture detection. Commun Biol 4:130. doi:10.1038/s42003-021-01654-9

Segalin C, Williams J, Karigo T, Hui M, Zelikowsky M, Sun JJ, Perona P, Anderson DJ, Kennedy A. 2021. The Mouse Action Recognition System (MARS) software pipeline for automated analysis of social behaviors in mice. Elife 10. doi:10.7554/eLife.63720

Shimada SG, LaMotte RH. 2008. Behavioral differentiation between itch and pain in mouse. Pain 139:681–687. doi:10.1016/j.pain.2008.08.002

Simpson EH, Akam T, Patriarchi T, Blanco-Pozo M, Burgeno LM, Mohebi A, Cragg SJ, Walton ME. 2024. Lights, fiber, action! A primer on in vivo fiber photometry. Neuron 112:718–739. doi:10.1016/j.neuron.2023.11.016

Smolinsky AN, Bergner CL, LaPorte JL, Kalueff AV. 2009. Analysis of grooming behavior and its utility in studying animal stress, anxiety, and depressionMood and Anxiety Related Phenotypes in Mice, Neuromethods. Totowa, NJ: Humana Press. pp. 21–36. doi:10.1007/978-1-60761-303-9_2

Sun G, Lyu C, Cai R, Yu C, Sun H, Schriver KE, Gao L, Li X. 2021. DeepBhvTracking: A novel behavior tracking method for laboratory animals based on deep learning. Front Behav Neurosci 15:750894. doi:10.3389/fnbeh.2021.750894

Vasanthi P, Mohan L. 2024. Efficient YOLOv8 algorithm for extreme small-scale object detection. Digit Signal Process 154:104682. doi:10.1016/j.dsp.2024.104682

Wen N, Guo R, Ma D, Ye X, He B. 2022. AIoU: Adaptive bounding box regression for accurate oriented object detection. Int J Intell Syst 37:748–769. doi:10.1002/int.22646

Whoriskey K, Auger-Méthé M, Albertsen CM, Whoriskey FG, Binder TR, Krueger CC, Mills Flemming J. 2017. A hidden Markov movement model for rapidly identifying behavioral states from animal tracks. Ecol Evol 7:2112–2121. doi:10.1002/ece3.2795

Wimalasena NK, Milner G, Silva R, Vuong C, Zhang Z, Bautista DM, Woolf CJ. 2021. Dissecting the precise nature of itch-evoked scratching. Neuron 109:3075–3087.e2. doi:10.1016/j.neuron.2021.07.020

Xu Z, Wang T, Skidmore AK, Lamprey R, Ngene S. 2025. Bounding box versus point annotation: The impact on deep learning performance for animal detection in aerial images. ISPRS J Photogramm Remote Sens 222:99–111. doi:10.1016/j.isprsjprs.2025.02.017

Yu H, Xiong J, Ye AY, Cranfill SL, Cannonier T, Gautam M, Zhang M, Bilal R, Park J-E, Xue Y, Polam V, Vujovic Z, Dai D, Ong W, Ip J, Hsieh A, Mimouni N, Lozada A, Sosale M, Ahn A, Ma M, Ding L, Arsuaga J, Luo W. 2022. Scratch-AID, a deep learning-based system for automatic detection of mouse scratching behavior with high accuracy. Elife 11. doi:10.7554/eLife.84042

Zhao Z-Q, Zheng P, Xu S-T, Wu X. 2019. Object detection with deep learning: A review. IEEE Trans Neural Netw Learn Syst 30:3212–3232. doi:10.1109/TNNLS.2018.2876865

